# ATF4 programs proline-dependent immune evasion in β-Catenin-driven hepatocellular carcinoma

**DOI:** 10.64898/2026.05.12.724215

**Authors:** Stefany Infante, Eva Santa Maria, Alex Finnemore, Sergio Barace, Angel Martinez-Montes, Francesco Damone, Sara Arcelus, Guillermo Garcia-Porrero, Nastaransadat Hosseini, Eduardo Miraval, Carlos E. de Andrea, Diana Llopiz, Maria Reig, Yaron Finkelstein, Bruno Sangro, Pablo Sarobe, Puri Fortes, Amaia Uriz, Juan Bayo, Josepmaria Argemi

**Affiliations:** Program of DNA and RNA Therapy, CIMA Universidad de Navarra, Cancer Center Clínica Universidad de Navarra (CCUN), Pamplona, Spain; Facultad de Medicina Humana. Universidad de Piura.; Centro de Investigacion Biomedica en Red de Enfermedades Hepaticas y Digestivas (CIBER-EHD), Madrid, Spain; Facultad de Medicina. Universidad de Navarra.; Program of Immunology and Immunotherapy, CIMA Universidad de Navarra, Cancer Center Clínica Universidad de Navarra (CCUN), Pamplona, Spain; Pathology Department. Northern Health and Social Care Trust. Antrim. Northern Ireland. United Kingdom; Pathology Department. Clinica Universidad de Navarra, Pamplona, Spain; Navarra Institute for Health Research (IDISNA), Pamplona, Spain; Barcelona Clinic Liver Cancer Group (BCLC), Barcelona, Spain; Departments of Pediatrics, Pharmacology and Toxicology, University of Toronto, The Hospital for Sick Children (SickKids), Toronto, Canada; Liver Unit and HPB Oncology Area, Cancer Center Clínica Universidad de Navarra (CCUN), Pamplona o Madrid, Spain; Gene Therapy Laboratory. Instituto de Investigaciones en Medicina Traslacional. CONICET Universidad Austral. Buenos Aires. Argentina; Division of Gastroenterology Hepatology and Nutrition. University of Pittsburgh. Pittsburgh. PA. USA

**Keywords:** Hepatocellular carcinoma, ATF4, Beta-catenin, Integrated stress response, Proline biosynthesis, Immune evasion, Tumor microenvironment, Macrophage polarization, Immunotherapy resistance, Metabolic-immune crosstalk

## Abstract

**Background & Aims:** Hepatocellular carcinoma (HCC) frequently exhibits resistance to immune checkpoint inhibitors (ICIs), particularly in β -catenin-driven tumors characterized by immune exclusion. While the Unfolded Protein Response (UPR) and the Integrated Stress Responses (ISR) enable tumor adaptation to metabolic stress their role in shaping tumor immunogenicity remains incompletely understood. We investigated whether ATF4, a central effector of the integrated stress response, couples metabolic reprogramming to suppression of anti-tumor immunity in HCC.

**Methods:** We combined transcriptomic analyses across three independent human HCC cohorts with mechanistic studies using an immunotherapy-resistant MYC/β-catenin-driven murine HCC model. We integrated CRISPR/Cas9-mediated deletion of *Atf4* with RNA-sequencing and targeted metabolomics. The impact of tumor-derived metabolites on macrophage differentiation and polarization was evaluated using primary bone marrow-derived cells. Therapeutic responses were evaluated in orthotopic and subcutaneous models treated with anti-PD-1 and anti-VEGFA.

**Results:** ATF4 and XBP1 transcriptional signatures are selectively enriched in human HCC and associate with poor prognosis, vascular invasion, and an immunosuppressive myeloid-enriched tumor microenvironment. Genetic ablation of *Atf4* markedly suppressed tumor growth in immunocompetent but not immunodeficient hosts, establishing a requirement for immune-mediated tumor control. Mechanistically, *Atf4* loss downregulated Aldh18a1 and disrupted proline biosynthesis, resulting in extracellular proline depletion. This proline-deficient environment abrogated monocyte-to-macrophage differentiation and decreased M2 polarization, thereby reshaping the tumor microenvironment toward enhanced T cell infiltration and activation. Functionally, Atf4-deficient tumors exhibited restored sensitivity to anti-PD-1 monotherapy and showed pronounced responses to combined anti-PD-1/anti-VEGFA treatment in aggressive orthotopic models.

**Conclusion:** ATF4 programs a proline-dependent metabolic axis that sustains macrophage-mediated immunosuppression and immune evasion in β-catenin-driven HCC. Disruption of this pathway converts immune-excluded tumors into T cell-inflamed states and restores responsiveness to immunotherapy. By governing proline homeostasis and macrophage-mediated immunosuppression, ATF4 is a key metabolic checkpoint for immune evasion, linking stress adaptation to immune escape and a candidate therapeutic target in HCC.

**Impact and implications:** We identify ATF4 as a crucial metabolic-immune orchestrator that sustains myeloid-driven immune evasion in β-catenin-dependent HCC through proline-dependent circuitry. Disrupting the ATF4-proline axis converts immune-desert tumors into T cell-inflamed lesions by blocking macrophage differentiation, thereby sensitizing tumors to immune checkpoint therapy. This work positions ATF4 as a tractable therapeutic target to overcome immunotherapy resistance in HCC.

**Graphical abstract:** 

**Highlights:**

- ATF4 orchestrates an immunosuppressive tumor microenvironment in HCC by coupling metabolic stress adaptation to immune evasion.
- Ablation of ATF4 disrupts proline biosynthesis, leading to a marked depletion of extracellular proline.
- Cancer cell-derived proline availability contributes to macrophage differentiation and M2 polarization; its loss restores T cell-mediated anti-tumor surveillance and sensitizes beta-catenin-driven HCC to immune checkpoint blockade.

## Introduction

Hepatocellular carcinoma (HCC) is a leading cause of cancer-related mortality worldwide (1). Despite advancements in immune checkpoint inhibitors (ICIs), efficacy is often limited by intrinsic or acquired resistance (2–4). To survive the hostile tumor microenvironment (TME), characterized by hypoxia and nutrient scarcity, HCC cells engage adaptive stress pathways, primarily the Unfolded Protein Response (UPR) and the Integrated Stress Response (ISR) (5–10).

The ISR centers on eIF2α phosphorylation by stress-sensing kinases (GCN2, PERK, PKR, HRI), leading to selective translation of Activating Transcription Factor 4 (ATF4) (11). While ATF4 maintains homeostasis in healthy cells (11, 12), in cancer it promotes chronic adaptation and survival (13). In the liver, ATF4 exhibits a dual role: during chronic inflammation, it can inhibit carcinogenesis by preventing ferroptosis (14), yet uncontrolled activation of its effector CHOP can exacerbate inflammation (15, 16). In established HCC, ATF4 drives progression (17) and confers resistance to cisplatin and tyrosine kinase inhibitors (18, 19). Beyond the ISR, ATF4 integrates one of the three UPR arms-alongside ATF6 and IRE1A/XBP1-to shape a landscape favoring tumor growth and immune evasion (20, 21). Notably, ATF4 was recently linked to immune evasion in KRAS-driven lung and pancreatic cancers via LCN2-mediated macrophage recruitment (22). However, the specific impact of the UPR/ISR on HCC immunity remains poorly defined.

In this study, we integrated transcriptomic analyses from human HCC cohorts with functional studies in a Beta-catenin-driven, immunotherapy-resistant HCC model. We demonstrate that, unlike ATF6 or XBP1, ATF4 activation defines a transcriptional program associated with poor prognosis and a macrophage-enriched, immunosuppressive TME across HCC etiologies. Genetic deletion of *Atf4* reduced tumor growth in immunocompetent, but not immunodeficient, models and restored sensitivity to ICIs. Mechanistically, *Atf4* loss disrupted proline biosynthesis, leading to defective monocyte-to-macrophage differentiation and impaired M2 polarization.

## Materials and Methods

### Patients and cohorts and clinical data

Bulk RNA-seq data and clinical annotations were obtained from three independent HCC cohorts: OEP000321 (OEP), The Cancer Genome Atlas Liver Hepatocellular Carcinoma (TCGA-LIHC, TCGA), and the International Cancer Genome Consortium Japan Liver Cancer-RIKEN (ICGC-LIRI-JP, LIRI). Expression matrices were downloaded as CPM values from the HCCDB database (HCCDB25, HCCDB15, and HCCDB18, respectively). For TCGA, only samples with confirmed hepatocellular carcinoma histology (NOS, spindle cell variant, and clear cell type) were retained. Tumor and adjacent non-tumor samples were classified using sample metadata. Clinical variables-including overall survival, BCLC stage, TNM stage, AFP levels, etiology (HBV, HCV, alcohol), vascular invasion, fibrosis score, and mutational status (*CTNNB1*, *TP53*)-were curated from the corresponding clinical files of each cohort. Gene symbols were harmonized to HGNC-approved nomenclature across cohorts.

### Generation and refinement of UPR/ISR transcriptional signatures

Transcriptional signatures for ATF4, ATF6, and XBP1 were curated and refined using a graph-based co-regulation approach as previously described (23). Candidate target gene sets for each transcription factor were assembled from the literature and processed through the *gsadapt* pipeline (https://github.com/unav-hcclab/gsadapt.git), which identifies co-regulated gene clusters via community detection. *gsadapt* was applied independently to tumor samples in each cohort to select modules enriched relative to non-tumor tissue.

For cross-cohort robustness, pairwise Pearson correlations of ssGSEA scores were computed per cohort using the *Hmisc* R package (v5.1). A signature co-occurrence network was constructed retaining pairs with correlation >0.6 across all three cohorts, and community detection was performed using the walktrap algorithm (*igraph*, v1.5.1).

### Generation of UPR ko cell lines

Stable knockout cell lines for the UPR transcription factors (UPR TF): XBP1, ATF4, ATF6 were generated using CRISPR/Cas9-mediated genome editing. Single guide RNAs (sgRNAs) were designed using Benchling (www.benchling.com) targeting early coding exons. For each gene, three independent sgRNAs were selected based on predicted on-target efficiency and minimal off-target activity. Tested and finally selected sgRNA sequences, according to the knockout efficiency, are provided in **Supplementary Table S1**. A non-targeting control guide (NTC) was used to generate unedited control cancer cells. Complementary oligonucleotides encoding each sgRNA were annealed and cloned into the BbsI restriction sites of pSpCas9(BB)-2A-Puro (PX459) (Addgene #62988), which co-expresses S. pyogenes Cas9 and a puromycin resistance cassette. Cells were transfected with PX459-sgRNA constructs using Lipofectamine 2000 (Thermo Fisher Scientific) according to the manufacturer’s instructions. 72 h post-transfection, cells were subjected to puromycin selection (1.75 µg/mL) for 2 weeks. Pool of surviving cells were functionally validated performing induction of endoplasmic reticulum stress by treating with tunicamycin (1 µg/ml) or amino acid deprivation during 8 hours. Loss of XBP1, ATF4, ATF6 or activity was confirmed by assessing canonical downstream target gene expression by RT-PCR (see oligonucleotide sequences in **Supplementary Table S2)** and protein levels (see list of antibodies **in Supplementary S3**), under normal and stress conditions.

### Orthotopic and subcutaneous models of HCC

Female C57BL/6J mice (7□weeks old) were obtained from Envigo. They were maintained in pathogen-free conditions and treated according to the EU Directive 2010/63/EU on the protection of animals used for scientific purposes, under a protocol approved by the Animal Research Ethics Committee of the University of Navarra (protocol 095-20). For the orthotopic HCC tumor model, 5 × 10^4^ MycBC cells (NTC and Atf4ko) were inoculated intra-hepatically. Tumor size was measured by ultrasound on day 11 post-inoculation to ensure each group had the same median tumor size baseline. Anti-PD-1 (BE0146, 100 ug) and anti-VEGFA (BE139337, 200 ug) antibodies were administered intraperitoneally (i.p.) every three days. An IgG isotype control (BE0090, 300 µg) was used in control mice. All antibodies used in vivo were from Bioxcell.

For the subcutaneous model, 1 × 10□ HCC cells were inoculated subcutaneously in the right flank of the animal. Treatment commenced on day 8 post-inoculation. Anti-PD-1 antibody and isotype control (100 µg each) were administered i.p. in 3 doses every three days. Tumor size was monitored twice per week with a caliper and volume was calculated with the following formula V = (a^2^ x b) / 2, where “a” is the smaller diameter and “b” the greatest diameter. Animals that developed a tumor of 1700 mm^3^ were sacrificed for ethical reasons.

To assess the role of the immune system in tumor growth, 1 × 10□ HCC cells were inoculated subcutaneously in *Rag2*^-/-^,*Il2r*^-/-^ mice (Jackson Laboratories, #038282) and tumor growth was monitored. In experiments of lymphocyte depletion, 1 × 10□ HCC cells were inoculated subcutaneously in 7□weeks old female C57BL/6J mice. After an initial high dose of depleting antibody in day -2 (200 ug/mouse), repeated doses at days 0, 3, 6, 9, 12 and 15 (100 ug/mouse). Tumor growth was monitored and when the tumors reached 1700 mm^3^ were sacrificed for ethical reasons. Anti-IgG (BE0090), Anti-CD4 (BE003-1) and anti-CD8 (BE0223), were all obtained from Bioxcell.

### In vitro cultures of bone marrow-derived macrophages

Mouse bone marrow macrophages were isolated from both femur and tibia of female 8-10 week old C57BL6J mice. Syringe-pushed medium was used to obtain a clean cylinder of bone marrow which was subsequently disaggregated in RPMI-1640 GlutaMAX medium (GIBCO-#61870-010). Sample was then subjected to erythrocyte lysis, quantified by trypan blue exclusion and cultured at a density of 1 x 10□ cells per well in a 12-well plate in RPMI medium containing 10% heat-inactivated fetal bovine serum (FBS), 1% penicillin/streptomycin and 0.1% β-mercaptoetanol.

For macrophage differentiation studies, 6 x 10□ cancer cells were cultured for 24 h in 12 ml of DMEM. The medium was then renewed and 8h later the supernatant was collected for its use in macrophage differentiation protocol. Bone marrow precursors were treated with cancer cell supernatant supplemented with 100 ng/ml of M-CSF (Inmunotools-#12343117). Media was renewed on day 3, at which point all non-adherent cells were removed. On day 6, macrophages were harvested and analyze by flow cytometry.

For polarization studies, macrophages were differentiated with 100 ng/mL M-CSF. On day 6, tumor cell supernatants were added along with M1-induction cocktail, including 50 ng/mL LPS (SIGMA- L2637-5MG) and 10 ng/mL IFNγ (Myltenyi- 130-105-774) or with 5 ng/mL of M2-induction factor IL-4 (Inmunotools-#2340045). After 24 h, cells were harvested for flow cytometry analysis.

## Results

### ATF4 and XBP1 signatures are increased specifically in HCC cancer cells regardless the etiology and the degree of the background liver damage

To understand the relevance of the stress response on tumor growth, we first conducted a comprehensive bioinformatic analysis of the unfolded protein response (UPR) and integrated stress response (ISR) mainstream transcription factor programs (**Figure 1A**). We curated experimentally validated mouse ChIP-seq datasets to identify bona-fide targets of Atf4, Xbp1 and Atf6. We then adapted the resulting gene sets to human transcriptomes using bulk RNA-seq data from three large independent HCC cohorts: a Chinese cohort (OEP000321) with 158 tumor and non-tumor pairs, the TCGA-LIHC cohort, with 350 tumor and 49 non tumor samples, and the Japanese ICGC-LIRI-JP cohort including 211 tumor and 175 non-tumor samples (**Figure 1B**), using a pipeline developed by our group in a previous work (23). By means of graph association based on single gene co-expression, 22 signatures were initially derived: 6 for ATF4, 8 for XBP1 and 8 for ATF6. These gene sets were further refined using signature co-expression networks, yielding 5 independent TF signatures, namely ATF4 (42 genes), XBP1-1 (175 genes), XBP1-2 (176 genes), ATF6-1 (88 genes) and ATF6-2 (46 genes) (**Figure 1B and Supplementary Figure 1A-C**). The resulting signatures were mostly independent with XBP1-1 and 2 signatures sharing only 28 genes and ATF6-1 and 2 sharing 15 genes, while only a very small fraction of genes were shared among the remaining signatures (**Supplementary Figure 1D).**

**Figure. 1.**
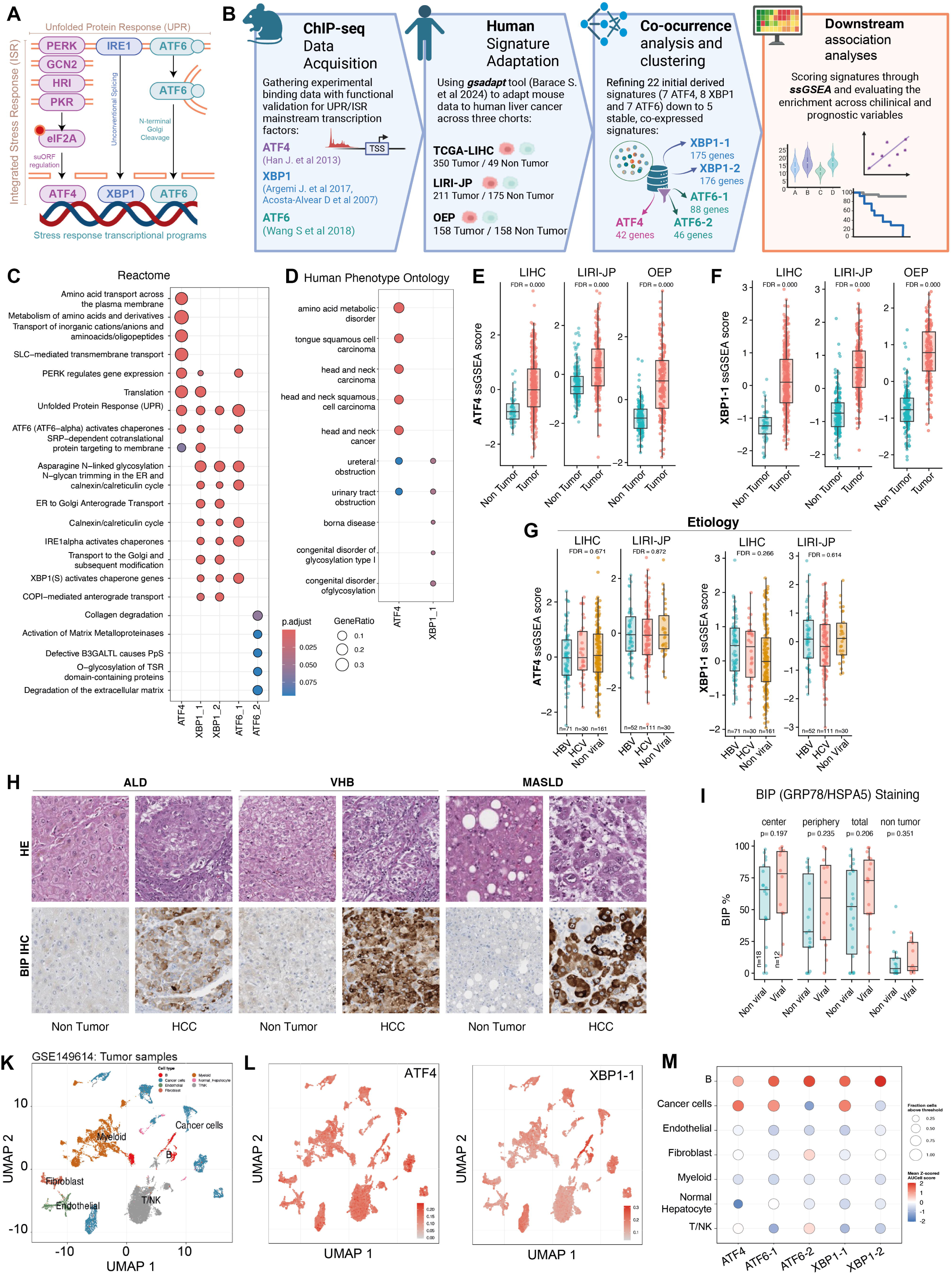
ATF4 and XBP1 signatures are increased specifically in HCC cancer cells regardless of the etiology and the degree of the background liver disease. (A) Unfolded protein response (UPR) and integrated stress response (ISR) is initiated in the ER lumen by three transmembrane sensors, PERK, IRE1 and ATF6 and their downstream transcriptional output is driven by three transcription factors: ATF4, XBP1 and ATF6 (B) Schema of the bioinformatic pipeline to define consistent ATF4, ATF6 and XBP1 signatures adapting ChipSeq data to the TCGA-LIHC, LIRI-JP-OEP cohorts of HCC, using gsadapt pipeline, deriving 5 highly co-expressed signatures (C-D) gene set enrichment analysis using Reactome (C) and Human Phenotype Ontology (D). (E-F) Signature scores of ATF4 (E) and XBP1-1 (F) using ssGSEA in the LIHC, LIRI-JP and OEP cohorts comparing non tumoral tissues vs tumors. (G) Signature scores of ATF4 and XBP1-1, splitting the LIHC population by the etiology of liver disease. (H) Immunohistochemistry of HSPA5 (also named GRP78 or BIP) in HCC and non-tumoral tissues from patients with different etiologies. (I) Quantification of H considering different areas of the tumor area. (J-L) Single cell sequencing analyses of ATF4, ATF6 and XBP1 signatures using AUCell Scores of a published dataset including 10 HBV-associated HCCs (GSE149614). (J) UMAP depicting the two principal components driving the different expression in the samples using randomly selected 20,000 cells from HCC samples, and colored according to the cell type. (K) UMAP colored by AUCell scores from ATF4 and XBP1-1 signatures according to the normalized intensity of the expression. (L) Mean expression and % of cells expressing the signatures of ATF4, ATF6 and XBP1, splitting the cell population by cell type.

The enrichment analysis using the Reactome collections confirmed the predicted role of ATF4 in amino acid metabolism and transport, protein translation and folding, including the UPR (**Figure 1C**). XBP1-1 signature revealed a predominant role in UPR, protein translation trafficking, glycosylation and folding and ER to golgi transport. XBP1-2 genes were related to a very similar functional profile than XBP1-1, but the latter uniquely include protein translation and the regulation of PERK and ATF6 related responses. ATF6-1enriched functions were very similar to XBP1-1 and 2, but surprisingly, despite its well-known activation cleavage by Golgi proteases, did not include any of the three pathways related ER-to-Golgi transport, present in XBP1 signatures’ enrichment. Finally, ATF6-2 signature functional enrichment had nothing in common with ATF4, XBP1 or ATF6-1 related pathways: only a poor enrichment of a few extracellular matrix metabolism gene sets could be enriched using this signature (**Figure 1C**). When interrogating the Human Phenotype Ontology gene sets, ATF4 signature was found enriched in amino acid metabolic disorder and different epithelial cancer gene sets, while XBP1-1 was found significantly associated to disorders of glycosylation (**Figure 1D**). These data indicate that the approach of generating transcriptional signatures from Chip-Seq assessed target genes yielded a coherent functional enrichment results. Complementary enrichment analyses using KEGG pathways (**Supplementary Figure 1E**) and Gene ontology collections, including Biological Process (**Supplementary Figure 1F**), Cellular Component (**Supplementary Figure 1G**), and Molecular Function (**Supplementary Figure 1H**), confirmed that ATF4, XBP1 and ATF6-1 share transcriptional outcomes related to protein translation and folding, while having specific functional enrichments. For example, signatures related amino acid metabolism and transport were specifically associated only with ATF4, while ER-to-Golgi and N-glycan biosynthesis were typical of XBP1 signatures. Interestingly, ATF6-1 signature was associated with Antigen processing and presentation, although with borderline significance (**Supplementary Figure 1E**).

We then explore the predicted activation of ATF4, XBP1 and ATF6 in HCC. ATF4 (**Figure 1E**) and XBP1-1 (**Figure 1F**) signatures were significantly increase in tumors when compared to non-tumoral tissues in all three cohorts explored. ATF6-1 signature was increased in tumors of two of the cohorts. In contrast, the XBP1-2 and ATF6-2 signature scores were not increased in tumors (**Supplementary Figure 2A-C**). These data suggest that ATF4 is probably associated with cancer cell metabolism or with the tumor environment, while for XBP1 only one of the two signatures derived could be involved in cancer. We thus aimed at studying the potential association of the UPR signatures with either TP53 or BC mutations the two most common driver genes in HCC (24). When analyzing the TCGA-LIHC cohort, an increase enrichment of ATF4 and XBP1-1 signature with the presence of TP53 missense or truncated mutations was found, an association that could not be validated in ICGC-LIRI-JP or OEP cohorts (**Supplementary Figure 2D**). Neither XBP1-2 nor ATF6 signatures were associated with the presence of TP53 mutation (**Supplementary Figure 2E-G**). We could not find consistent association of BC-mutation with the ATF4, XBP1-1 (**Supplementary Figure 2H**) or XBP1-2 and ATF6 (**Supplementary Figure 2I-K)** signatures, apart from a decrease of ATF6-2 signature in LIHC (**Supplementary Figure 2J**) and a decrease of the signature of XBP1-1 signature in OEP (**Supplementary Figure 2H**). HCC normally arises in the context of chronic liver disease, either viral (i.e. induced by chronic Hepatitis B and C virus infection) or non-viral (i.e. alcohol or metabolic-related). The different immune microenvironment and the hypothetical differential response to therapy according to the etiology have been a focus of attention in recent years (25). On the other hand, the UPR has been associated to fatty liver disease (26), Hepatitis B (27) and Hepatitis C (28) pathogenesis and disease progression. In LIHC and LIRI cohorts, none of the 5 signatures were consistently associated with the etiology of the liver disease (**Figure 1G, Supplementary Figure 3A**). ATF4, ATF6-2 and XBP1-2 tended to be higher in the HCC of obese patients in the TCGA-LIHC cohort (**Supplementary Figure 3B)**. To further confirm the lack of strong association with the etiology, we performed an IHC of Heat Shock Protein 5a or Glucose-regulated protein 78 (HSPA5, GRP78 or BIP), a well-known UPR target shared by the three branches of the UPR under stress conditions. We could detect increased cytoplasmic HSPA5 staining in HCC both from viral and non-viral etiology when compared to non-tumoral liver tissue (**Figure 1H-I**). HSPA5 was broadly positive in tumor cells, with no gross spatial differences, while non-tumoral cells, even considering metabolic liver or HBV chronic disease background had significantly low HSPA5 cytoplasmic abundance (**Figure 1I**). These data show that ATF4 and XBP1-1 signatures are enriched in HCC regardless the driver gene and the etiology of the background liver. The UPR transcriptional activity in the tumor was not associated with the presence in the background liver of inflammation (**Supplementary Figure 4A**) or fibrosis per Ishack score (**Supplementary Figure 4B**).

Finally, to understand the cellular source of the expression of ATF4, XBP1 and ATF6 signatures, we interrogated the UPR transcription factors’ signatures in a single cell experiment from 10 patients with HBV-associated HCC, previously published (29). The immune microenvironment was well represented in these samples and included Myeloid, Fibroblast, Endothelial, T/NK Cells, B Cells, Cancer cells and a few Hepatocytes (**Figure 1J**). The range of normalized expression per cell was lower for ATF4 signature when compared to XBP1-1 (**Figure 1K**), due to a particularly enhanced expression of the latter in B Lymphocytes. This aligns with the fact that XBP1 is essential for B cell differentiation to plasma cells and in general with the evidence of ER stress and UPR in professional immunoglobulin secretory cells (30). In this sense, it was not surprising to find that ATF4, XBP1-2 and ATF6 were also increased in B cells. Only in the case of ATF4 signature, the predicted activation was higher in Cancer cells than in B cells and any other cell type. ATF6-2 was also increased in fibroblasts and T/NK. Neither ATF6-2 nor XBP1-2 have increased scores in cancer cells (**Figure 1L**).

### ATF4 and XBP1 are associated to more aggressive HCC phenotype

We then explored ATF4 and XBP1-1, which are increased in HCC, in relation to tumor aggressiveness. Interestingly ATF4 and XBP1-1-inferred activity was increased in the proliferation class and decreased in Poly7 of Chiang transcriptomic subclass (31) (**Figure 2A**). ATF4 was increased in S1 and decreased in S3 Hoshida subclasses (32), while XBP1-1 was decreased in S3 Hoshida subclass (**Figure 2B**). No association with transcriptomic subclasses of Chiang or Hoshida was found for ATF6 or XBP1-2 signatures (**Supplementary Figure 5A-B**). Strikingly, ATF4 score was high in tumors with vascular invasion in both LIHC and LIRI cohorts, with a trend also for XBP1-1 (**Figure 2C-D**) which was not the case for the remaining signatures (**Supplementary Figure 5C-D)**. Accordingly, highly de-differentiated tumors in both cohorts had increased scores of ATF4 and XBP1-1, with a linear positive trend from well to poor differentiated (**Figure 2E**) a feature not seen with ATF6 or XBP1-2 signatures (**Supplementary Figure 5E-F**). These results suggest that ATF4 and XBP1-1-high tumors are highly proliferative, poorly differentiated and prone to invasion. Interrogating both signatures, unbiasedly, in all tumor and non-tumor samples from LIHC cohort, using a pseudotime analysis, we could confirm that high-ATF4/XBP1-1 tumors are at the most distant stage when compared with non-tumors, with an intermediate evolutive stage group, showing a linear inferred evolution of ATF4 and XBP1 activation from healthy liver to cancer development and progression (**Figure 2F and G**). Finally, we explored the potential prognostic impact of ATF4 and XBP1-signatures. In a very consistent fashion, patients with high levels of ATF4 signature had poorer overall survival in LIHC (**Figure 2H**), LIRI (**Figure 2I**) and OEP (**Figure 2J**). Similarly, patients with high XBP1-1 had worse prognosis in the three cohorts (**Figure 2K-M**). As a sign of specificity, the scores of none of the remaining UPR signatures (XBP1-2, ATF6-1 and 2) marked a group of patients with different prognosis (**Supplementary Fig 6A-C**).

**Figure 2.**
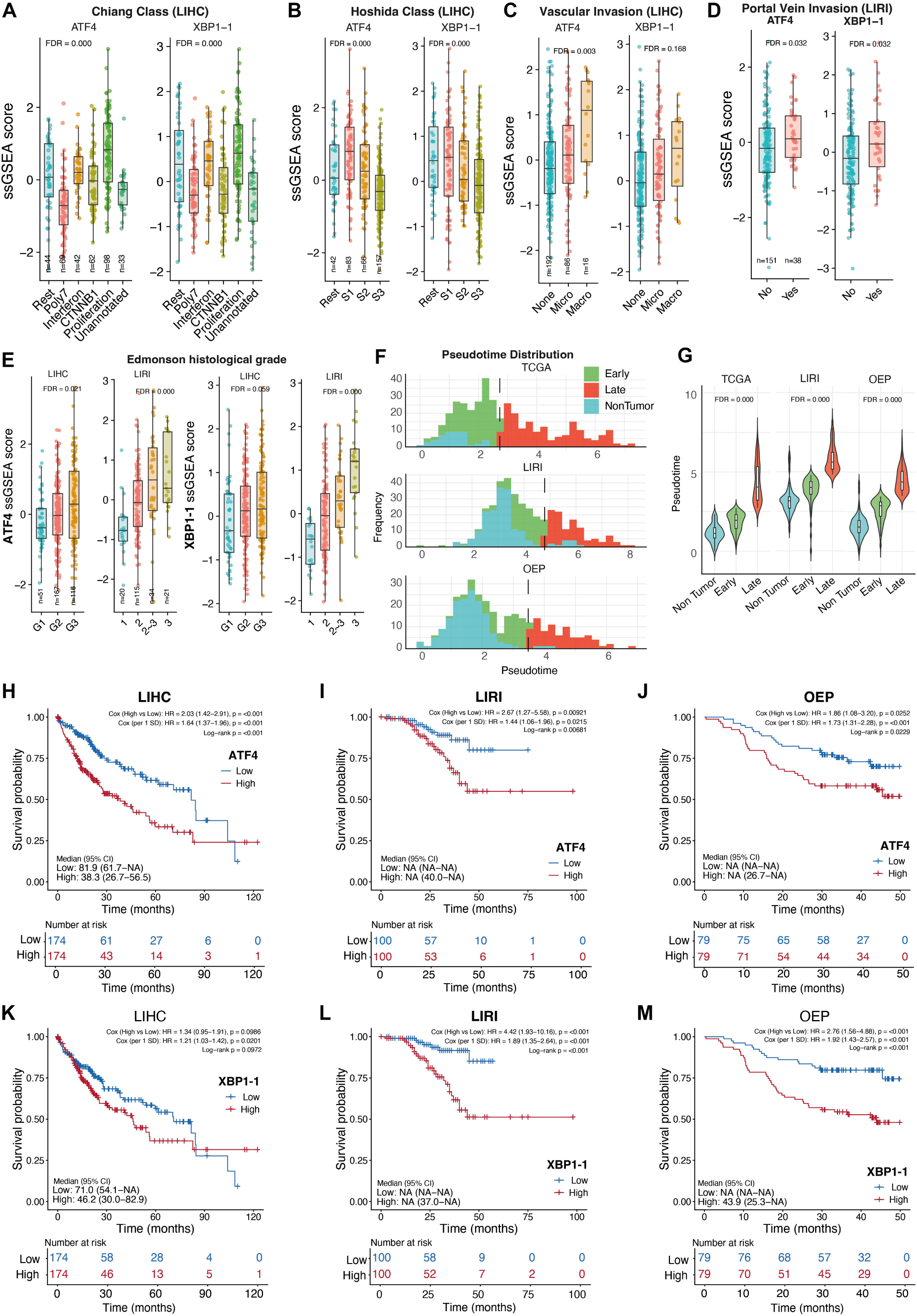
ATF4 and XBP1 are associated to more aggressive HCC phenotype. (A) Signature scores (ssGSEA) of ATF4 and XBP1-1 in LIHC cohort according to the Chiang transcriptomic subclasses (B) Signature scores (ssGSEA) of ATF4 and XBP1-1 in LIHC cohort according to the Hoshida transcriptomic subclasses. (C) Signature scores (ssGSEA) of ATF4 and XBP1-1 according to the presence of vascular invasion in the LIHC cohort. (D) Signature scores (ssGSEA) of ATF4 and XBP1-1 according to the presence of portal vein invasion in the LIRI cohort. (E) Signature scores (ssGSEA) of ATF4 and XBP1-1 according to the degree of differentiation per histological characterization according to Edmonton Scale (G1 well differentiated HCC; G2 moderate differentiation; G3 poorly differentiated HCC) in LIHC and LIRI cohorts. (F-G) Pseudotime analysis using ATF4 and XBP1-1 signatures, including the (F) histogram of frequences of samples with different pseudotime scores (non-tumor=light blue, early time HCC=green, late time HCC=red). (G) Violin plot of the pseudotime value, grouping by non-tumor, HCC early and HCC late. (H-J) Overall survival analysis of cohorts LIHC (H), LIRI (I) and OEP (J) splitting the population by high or low expression of ATF4 signature (ssGSEA). (K-M) Overall survival analysis of cohorts LIHC (K), LIRI (L) and OEP (M) splitting the population by high or low expression of XBP1-1 signature (ssGSEA).

Altogether, these data suggest that UPR/ISR signaling is tightly linked to aggressive tumor behavior in human HCC and should be further explored in functional studies.

### Atf4, Atf6, and Xbp1 knockout reveal limited effects on proliferation in vitro

Given the potential relevance of UPR/ISR transcription factor activity in patients with HCC, we next generated knockout (KO) cell lines for Xbp1, Atf4 and Atf6 using CRISPR/Cas9 in a murine HCC cell line that harbors MYC overexpression and β-catenin truncation (MycBC) and is partially resistant to immune checkpoint blockade, thanks to the presence of a composite ovoalbumin antigen in frame with luciferase gene. This unique model creates an immune suppressed environment resembling human beta-catenin-driven HCC (33). In the original paper, tumors directly originated from hydrodynamic tail vein injection were consistently immune excluded. In our experiments we derived a single clone that produced reproducible tumors in vivo that were immune suppressed, meaning with a few lymphocyte but important macrophage infiltration. The efficiency of the knockout was validated under ER stress conditions using amino acid deprivation or tunicamycin (Tm), confirming the efficient loss of function of Atf4 (**Supplementary Figure 7A-B**), Atf6 (**Supplementary Figure 7C-D**) and Xbp1 (**Supplementary Figure 7E-F**) when compared to cells treated with non-targeted sgRNA control guides (NTC). We next assessed possible compensatory activation among the three branches of the UPR. Interestingly, while Xbp1ko cells displayed mild upregulation of Atf4 and Atf6 under stress conditions and Atf4ko cells showed increased expression of Xbp1 and Atf6, the Atf6ko cells showed minimal compensation (**Supplementary Figure 7G**).

Despite these molecular alterations, all KO lines exhibited same proliferation under basal conditions (**Supplementary Figure 7H**). Furthermore, viability assays revealed no major differences in cell survival across genotypes when cultured under amino acid deprivation (**Supplementary Figure 7I**) or Tm (**Supplementary Figure 7J)**. Amino acid deprivation and tunicamycin exposure led to a sharp reduction in cell viability across all lines. ER stress responses drive life-death decisions in normal cells, and some of the output of the three transcription factors include both pro-survival and pro-apoptotic targets (34). Amino acid deprivation strongly induced apoptosis in all lines, though Atf4ko cells showed partial resistance. Likewise, when treated with Tm, the increase in apoptosis was lower in Atf4ko. In this second condition Xbp1 and Atf6-defficient cells experienced higher degree of apoptosis (**Supplementary Figure 7K**). This could indicate the lack of pro-apoptotic Ddit3 (or Chop) expression, a direct Atf4 target, impairing cell death pathways upon unsolved stress (35). We then sought to understand the impact of Atf4, Atf6, and Xbp1 on the cell cycle of MycBC cells. Tm-induced deficient glycosylation led to cell cycle arrest in G0-G1 phase in all conditions. In both baseline and Tm-treated conditions Atf4ko exhibited an arrest in G2-M phase when compared to NTC while Xbp1-loss also led to cell stacking in G2-M when compared to NTC but only in non-stressed conditions (**Supplementary Figure 7L)**. We and others have described the role of Xbp1 in in regenerating liver under replicative stress (36). On the other hand, it is known the cooperation between Myc and Atf4 in cell proliferation (37).

### Atf4 but not Xbp1 or Atf6 deletion reduces in vivo tumor growth and reshapes the immune infiltrate landscape

To investigate the *in vivo* consequences of UPR/ISR transcription factor loss, we evaluated tumor growth in subcutaneous models. Following subcutaneous implantation of syngeneic MycBC cells in an immunocompetent mouse model (**Figure 3A**), the tumors were generated in all animals and were able to growth, indicating their immune-evasion capacity. We thus implanted NTC, along with Xbp1, Atf4 and Atf6 ko cells. Strikingly, despite minimal differences in proliferation previously seen in *in vitro* conditions, Atf4ko tumors exhibited markedly reduced growth when implanted into immunocompetent C57BL/6J mice. In contrast, Atf6-loss led to a non-significant increase in tumor size, and Xbp1 deletion did not affect tumor growth in vivo (**Figure 3B y C**).

**Figure 3.**
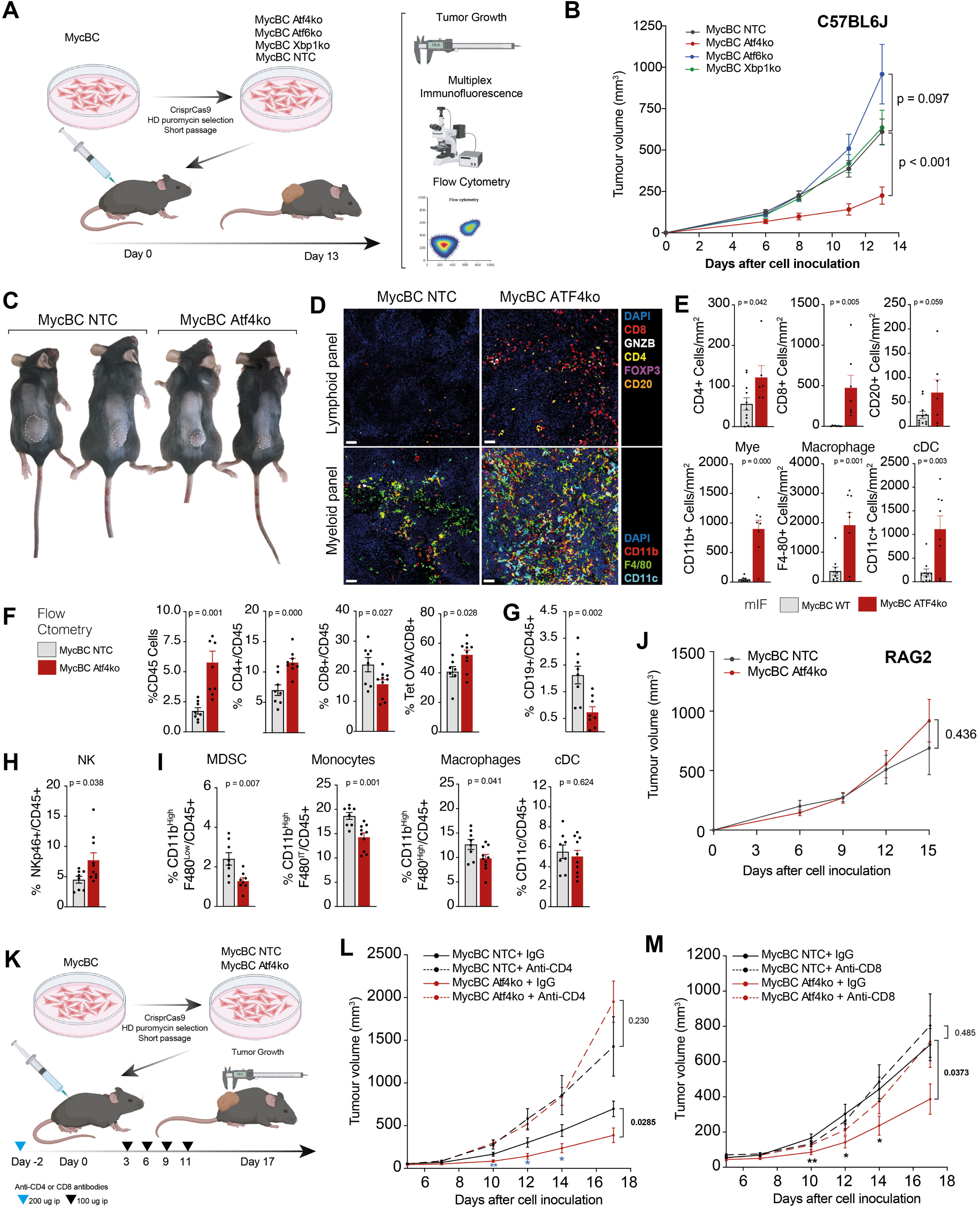
Atf4 but not Xbp1 or Atf6 deletion reduces in vivo tumor growth and reshapes the immune infiltrate landscape. (A) Schematic Immunocompetent C57BL/6J mouse models inoculated subcutaneously with UPR TF KO cell lines. At 13 days post inoculation, tumor growth immunofluorescence multiplex and flow cytometry was performed (B) Tumor growth measurements 13 days post cell inoculation. Data are shown as mean ± SEM. Statistical significance was assessed using two-way ANOVA with appropriate multiple comparison correction. (C) Representative images of tumor size in mice inoculated with MycBC NTC or Atf4ko cells. (D) Representative multiplex immunofluorescence images of MycBC NTC and Atf4ko tumor sections. Lymphoid panel: detection of CD8 (red), GZMB (white), CD4 (yellow), FOXP3 (magenta), and CD20 (orange), with DAPI (blue) as a nuclear marker. Myeloid panel: detection of CD11b (red), F4/80 (green), and CD11c (cyan), and DAPI (blue). (E) Quantification of positive cells per mm² for lymphoid (CD4^+^, CD8^+^, CD20^+^) and myeloid (CD11b^+^, F4/80^+^, CD11c^+^) populations in tumors. Data are presented as mean ± SEM, with each point representing an individual tumor. Statistical significance was assessed using Unpaired Student’s t test. (F-I) Flow cytometry analysis of immune-infiltrating populations in tumors. The percentage of lymphoid and myeloid cells is represented over the CD45^+^ population. Data are presented as mean ± SEM, with each point representing an individual tumor. Statistical significance was assessed using a two-tailed Unpaired Student’s t test. (J) Tumor growth curves of Immunodeficient RAG2-/- (*n*□=□8 mice/group). (K) Schematic overview of depletion of (L) CD4^⁺^ and (M) CD8^⁺^ T cell populations in Immunocompetent C57BL/6J mice (*n*□=□12 mice/group). Data are shown as mean ± SEM. Statistical significance was assessed using two-way ANOVA with multiple comparisons test and Uncorrected Fisher’s LSD.

To further investigate the tumor phenotype associated with Atf4 deficiency, we performed multiplex immunofluorescence on tumor sections. Atf4ko tumors exhibited increased global immune infiltration including CD4^⁺^ and CD8^⁺^ T cells, including granzyme B^⁺^ CD8^⁺^ cytotoxic T lymphocytes, as well as elevated numbers of dendritic cells, monocytes, and macrophages (**Figure 3D-E**). In order to reveal the relative abundance of each immune cell type and the presence, tumors were analyzed by flow cytometry. In Atf4-deleted tumors, an overall increase in CD45^⁺^ leukocyte infiltration was observed. Within the CD45^⁺^ compartment, there was a marked shift toward higher proportions of CD4^⁺^ T cells and tumor-specific CD8^⁺^ T cells (identified by OVA-tetramer staining), despite a relative reduction in total CD8^⁺^ T cell frequency (**Figure 3F**). In contrast, Atf4-deficient tumors showed decreased levels of CD19^⁺^ B cells (**Figure 3G**) and increased infiltration of NKp46^⁺^ natural killer cells (**Figure 3H**). Regarding the myeloid compartment, Atf4-deleted tumor infiltrates were lower in monocytes, macrophages and myeloid-derived suppressor cells (MDSCs) (**Figure 3I**). Overall, these results points towards a substantial reshaping of the immune infiltrate composition in Atf4ko tumors, consistent with a reduction in an immunosuppressive environment and an increased tumor-specific recognition. This could imply that Atf4 is directly involved in immune evasion. Since Atf4 could impair Myc-proliferative capacity regulating proteotoxic stress, we could not discard a partial immune system-independent effect of Atf4 deficiency in tumor growth. To test this hypothesis, we first implanted MycBc-NTC and Atf4ko cells into immunodeficient RAG2 mice. The differences in tumor size previously seen in immunocompetent mice, were completely lost (**Figure 3J**), indicating a complete dependency on T cells and/or NK cells of Atf4ko tumor rejection phenotype.

To functionally validate the immune relevance and the role of T cells in tumor control, we performed CD4^⁺^ and CD8^⁺^ T cell *in vivo* depletion experiments (**Figure 3K**). Depletion of either subset abrogated the growth delay of Atf4ko tumors, rendering them comparable to NTC tumors. Notably, CD4^⁺^ T cell depletion led to markedly accelerated tumor growth highlighting the critical role of CD4 T cells in the natural anti-tumor surveillance in this model (**Figure 3L**). CD8^⁺^ T cell depletion showed a similar pattern: while the effect of the depletion in the natural growth of MycBC tumors was less evident, the differences in tumor growth between NTC and Atf4ko tumors were abolished upon depletion (**Figure 3M**). These results suggest that Atf4 renders MycBC tumors less vulnerable to CD4/CD8 anti-tumor surveillance.

### Atf4 loss impairs amino acid usage leading to extra-cellular proline and aromatic amino acid depletion

Very recently, Atf4 in a Kras model of lung cancer has been shown to drive the expression of lipocalin 2 (Lcn2), promoting the infiltration of immunosuppressive interstitial macrophages and T cell exclusion (22). In endometrial cancer, Atf4 regulates Ccl2 to promote an immune suppressive environment (38). In our liver cancer MycBC model, the expression of Lcn2 and Ccl2 was negligible, excluding this mechanism as the explanation of the changes in the immune microenvironment. To understand the cancer-cell intrinsic mechanism behind the higher immune effector infiltration in MycBC tumors, we performed an RNA-sequencing of NTC and Atf4ko cells (**Figure 4A**). The deletion of Atf4 led to important changes, mostly towards the downregulation of several classical Atf4 programs, including the ISR and ER Stress responses, amino acid catabolism, synthesis and tRNA charging and mitophagy/hypoxia-related (**Figure 4B-C**). Among the hypoxia-related genes, Vegfa is known to be bound and regulated by Atf4 in cancer cells and has a pro-tumorigenic role by inducing angiogenesis and a first line target in systemics treatment for patients with HCC (39). Nevertheless, the strongest signal in our RNA sequencing data was related to amino acid metabolic, biosynthetic pathways, which have been related to potential immune suppressive mechanisms, not fully explored in HCC. Atf4-deficent tumor had a strong downregulation of rate limiting enzymes such as Aldh18a1, encoding P5CS (pyrroline-5-carboxylate synthetase), essential for the *de novo* proline biosynthesis, and Psph or phosphoserine phosphatase, essential for *de novo* serine biosynthesis and, together with mitochondrial Shmt2 and Mthfd2, are crucial for the one-carbon metabolism. Proline biosynthetic pathway is increased in Myc overexpressing tumors, and its blockade decreases tumor cell growth -including nucleotide synthesis- and energy production -including glycolytic activity- (40, 41) but the effect on immune evasion has not been described so far. Serine biosynthetic pathway (SBP) provides the one-carbon units required nucleotide, protein and phospholipid synthesis in highly proliferating tumor cells. More importantly, SBP induces an immunosuppressive phenotype by promoting M2-type macrophage polarization (42). Finally, Asns (asparagine synthetase) which produces asparagine from aspartate and glutamine and is one of the most canonical Atf4 targets is increased expressed in cancer. Loss of Asns forces cancer cells to depend entirely on extracellular asparagine. Importantly, Asns depletion in bladder cancer strongly limited tumor growth in a CD8+ T cell-dependent manner and boosted immunotherapy (43).

**Figure 4.**
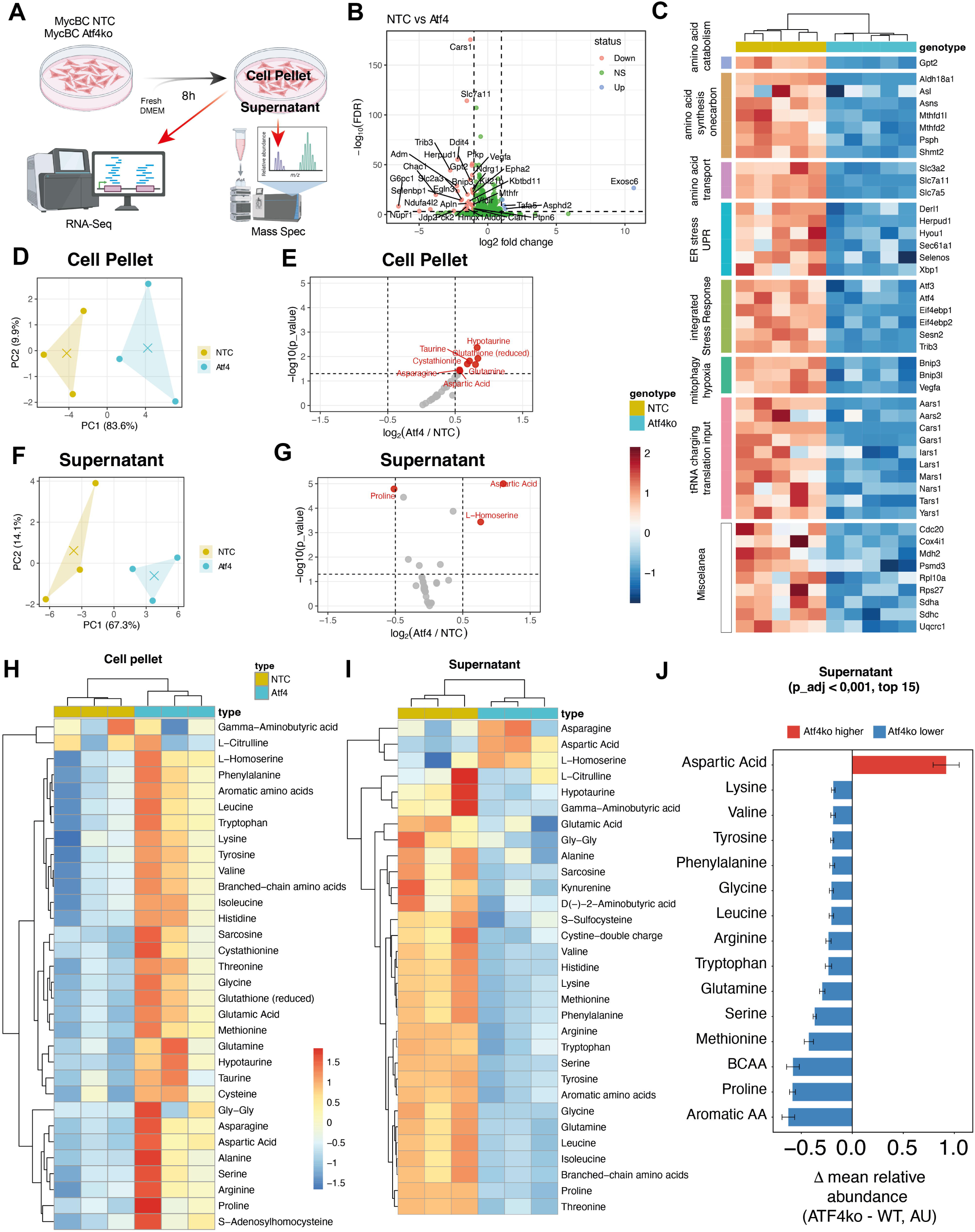
Atf4 loss impairs amino-acid usage leading to extracellular proline and aromatic amino acid depletion. (A) Schematic of the experimental workflow combining RNA-Seq and mass spectrometry analysis of cell extracts (Cell Pellet) and culture media (Supernatant) of MycBC NTC and Atf4ko cells after 8 hours of incubation. (B) Volcano plot and (C) RNAseq heat analysis. showing the transcriptional signature of genes involved in amino acid catabolism and transport, endoplasmic reticulum stress response (UPR), integrated stress response (ISR), and mitophagy. (D-G) Principal component analysis (PCA) and volcano plots of metabolomic data in MycBC NTC and ATF4Ko Cell pellet and supernatant. (H-I) clustering heat maps showing the relative abundance of metabolites in the cell and supernatant, (J) Mean relative abundance of the 15 most affected secretion of essential and branched-chain amino acids (BCAAs) in the absence of ATF4 in the supernatant (p < 0.001). Metabolomic data are presented as mean ± SEM.

We hypothesize that the impact of Atf4 in the usage and catabolism of amino acids could be linked to changes in the amino acid pool composition both inside and outside the cells, impacting in the tumor microenvironment. We thus performed targeted amino acid measurement of the intra- and extracellular compartments. Cell pellets and culture supernatants of Atf4-deleted and wt cells were analyzed through HPLC at 0 h and 8 h post-incubation (**Figure 4A**). The amino acid landscape both within cancer cells (**Figure 4D-E**) and in the extracellular medium (**Figure 4F-G**) was highly dependent on Atf4. Atf4 deletion caused broad reverse remodeling of amino-acid composition when compared the intra- versus the extracellular compartments. Notably, most amino acids accumulated in the intracellular compartment (**Figure 4E-H**) while in the cell supernatant most amino acid were relatively depleted. When using strict thresholds including the consideration of cell biomass (protein amount) and cell count, only proline was consistently depleted from the media (**Figure 4G**) while aspartate and L-homoserine were increased. When looking at the general trends comparing the two compartments, the results of amino acid quantification were less robust in cell pellets (**Figure 4H )** when compared to supernatants (**Figure 4I-J**). The depletion of amino acid in the supernatant was generalized with aspartate and l-homoserine being the exception (**Figure 4G-I**). Interestingly, proline, serine and methionine, the three amino acids related to Pshp and the SBP were found to be the most depleted in the supernatants of Atf4ko cells, while aspartic acid was highly accumulated, maybe reflecting the lack of catabolism derived from Asns deficiency (**Figure 4J )**.

### Cancer cell Atf4 governs distant macrophage differentiation by increasing the extracellular proline secretion

Cancer cells with proline synthesis deficiency and media proline depletion could lead to a complete shift of the metabolic components of the tumor microenvironment known to be important hallmarks of immune evasion. Two of them, namely the tumor-associated macrophages (TAM) and the cancer-associated fibroblasts (CAF) are known to be highly dependent on amino acid pool and/or on proline specifically. Amino-acid starvation, results in defective M2 polarization and enhanced M1 polarization (44) and proline supplementation can suppress M1 polarization by 50% in a model of radiation-induced inflammatory macrophage phenotype (45). Since Atf4ko tumors had a relative decreased monocyte and macrophage infiltration, we therefore explored the possibility that low proline environment could affect macrophage differentiation and/or polarization.

Primary mouse macrophages were obtained from bone marrow progenitors following a standard protocol. In the first 24h, the cells were treated with M-CSF and the treatment was repeated at day 3. At day 6 most of the cells have become M0 macrophages and are ready to be polarized into M1 or M2 using the appropriate stimuli (LPS/IFNg or IL4, respectively) (**Figure 5A**). To verify the dependency of monocyte-to-macrophage differentiation on proline levels, we first cultured bone marrow progenitors with complete RPMI media and proline-depleted RPMI. The removal of proline from the media abrogated the differentiation into macrophage almost completely (**Figure 5B**). DMEM-cultured cells reached a suboptimal level of differentiation (**Figure 5B**). Of note, one of the differences between DMEM and RPMI is the amino acid components: while DMEM, used in cancer cell cultures, is richer in aromatic amino acids (phenylalanine, tyrosine, and tryptophan), RPMI uniquely contains proline, aspartate, absent in DMEM (**Supplementary Table S5**). Cancer cells alter amino acid composition through import, export, and catabolism. This may explain the relative accumulation of aspartate in Atf4KO cells and increased proline levels in NTC controls (**Figure 4**). Next, supernatants from NTC and Atf4ko cells were used to replace monocyte media in the process of primary macrophage differentiation and polarization protocols. Cultured under supernatant from NTC cells, a decreased differentiation was observed when compared to fresh DMEM, indicating the capacity of tumors to modify macrophage differentiation by means of nutrient depletion. Interestingly, supernatant from Atf4 ko cells induced a significant decrease in macrophage differentiation when compared to cells that received supernatant from NTC cells. The addition of proline to the conditioned medium to Atf4-ko but not to NTC cells, restored macrophage differentiation (**Figure 5C**). This data suggests that proline depletion is an essential element defining the impact of Atf4ko supernatant on monocyte-to-macrophage differentiation.

**Figure 5.**
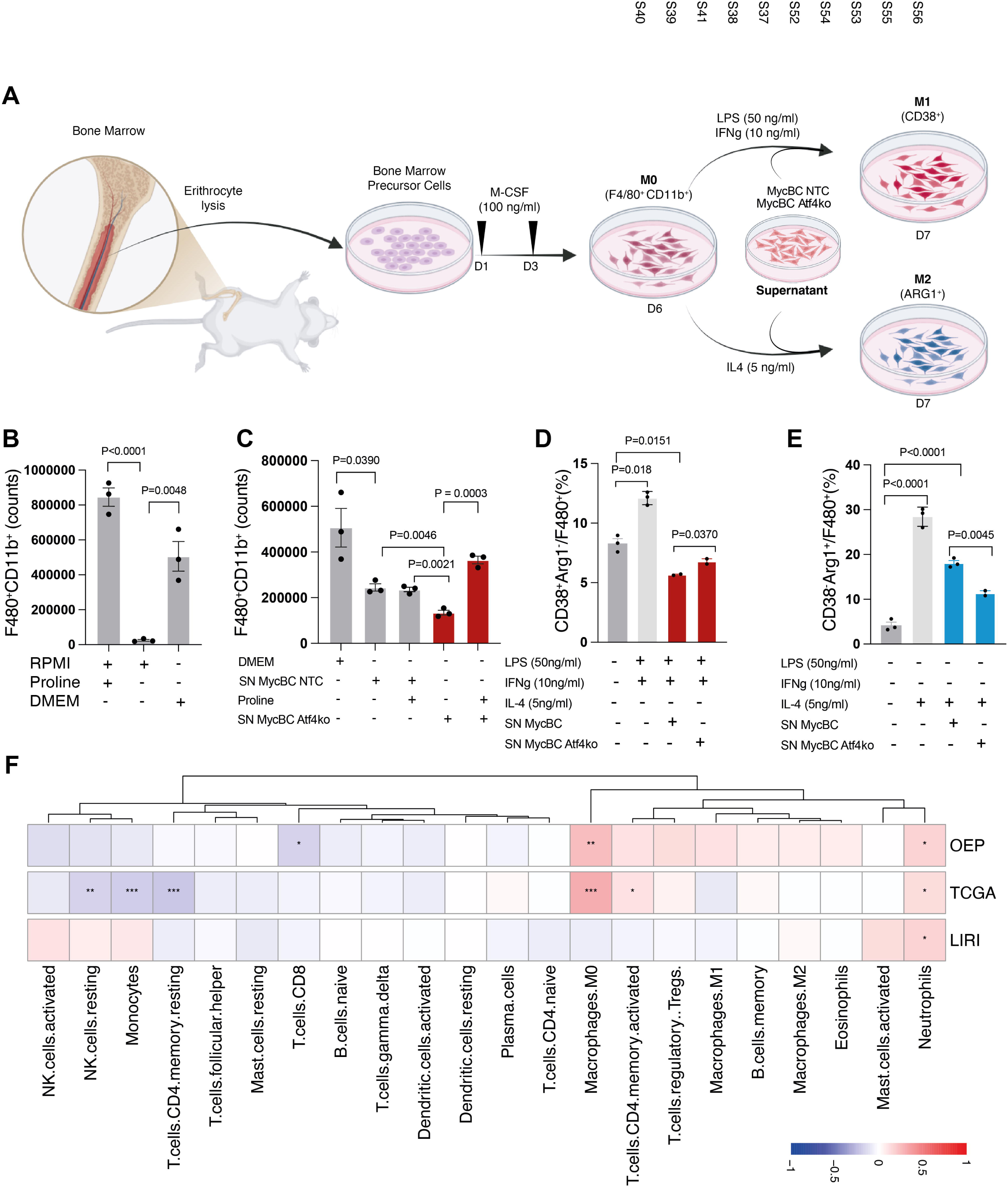
Cancer cell Atf4 governs macrophage differentiation by increasing the extracellular proline secretion. (A) Schematic representation of the bone marrow-derived macrophage (BMDM) differentiation protocol toward M0 (F4/80^+^ CD11b^+^), M1 (CD38^+^), or M2 (ARG1^+^) phenotypes, including exposure to MycBC NTC or Atf4ko cell supernatants (SN) and chemokine stimulation for M1 or M2. (B-C) Comparison of F4/80^+^ CD11b^+^ macrophages cell counts in different media conditions (RPMI, DMEM, and RPMI without proline) and proline supplementation in macrophages differentiation assays. (D) Percentage of M1 (CD38^+^ Arg1^-^) and (E) M2 (Arg1^+^ CD38^-^) in macrophages polarization assays by flow cytometry analysis. (F) immune cell composition of the LIHC, LIRI and OEP cohorts correlated with ATF4 signatures. Data are presented as mean ± SEM. Statistical significance was assessed using Unpaired Student’s t test.

Next, we sought to investigate if Atf4-derived proline depletion could also impact on macrophage polarization. Even though DMEM is not the ideal media for macrophage polarization, we could achieve an increase in M1 and M2 population upon treatment with LPS/IFNg and IL4, respectively When treated with NTC cell supernatant, the polarization of M0 macrophages to both M1 or M2 was impaired (**Figure 5D-E**). Interestingly, the supernatant of Atf4 induced a slightly less reduced M1 polarization (**Figure 5D**) and a more profound suppression of M2 phenotype (**Figure 5E**). To verify that the relationship between tumor Atf4 and macrophage differentiation is plausible in human HCC, we deconvoluted immune cell composition of the LIHC, LIRI and OEP cohorts and performed a correlation with ATF4 signature (**Figure 5F**). Strikingly, for LIHC and OEP cohorts -not for LIRI- there was a significant positive correlation of ATF4 signatures with Macrophage M0 inferred infiltration. Neutrophils were positively associated with ATF4 signature in all three cohorts. Resting NK, memory CD4 and monocytes were negatively associated with ATF4 signature in TCGA while CD8 T cells had the same trend, but only significant in OEP (**Figure 5F**). Collectively, these experiments show that Atf4 in cancer cells drives the differentiation and M2 polarization of macrophages, potentially explaining a more suppressive tumor microenvironment.

### Atf4 deletion restores immunotherapy sensitivity and correlates with immunosuppressive features in human HCC

All together, these observations pointed towards Atf4 as an orchestrator of the tumor-immune cross-talk by inducing a proline-driven macrophage differentiation, which, in turn, leads to the generation of a suppressive TME. In human HCC, macrophage infiltration is one of the main drivers of the primary resistance of patients with HCC treated with immunotherapy (46–48). We then hypothesize that Atf4 deficiency could impact immunotherapy regimens. We first performed an orthotopic model of HCC, by injecting MycBC cells in the median lobe of immunocompetent C57BL6J mice and treated them with twice weekly doses of murine anti-PD1 plus anti-VEGFA monoclonal antibodies, pathways currently targeted by one of the first line treatment combinations in HCC (**Figure 6A**). In this model, contrary to what was observed with subcutaneous tumors, NTC and Atf4-ko tumors had similar growth, indicating a more suppressive phenotype of orthotopic tumors (**Figure 6B-C**). Upon treatment with the combination, NTC tumors did not experience significant responses. Noteworthy, the growth Atf4-ko tumors treated with anti-PD1+anti-VEGFA immunotherapy was highly suppressed (**Figure 6B-C**), leading to important differences in survival (**Figure 6D**).

**Figure 6.**
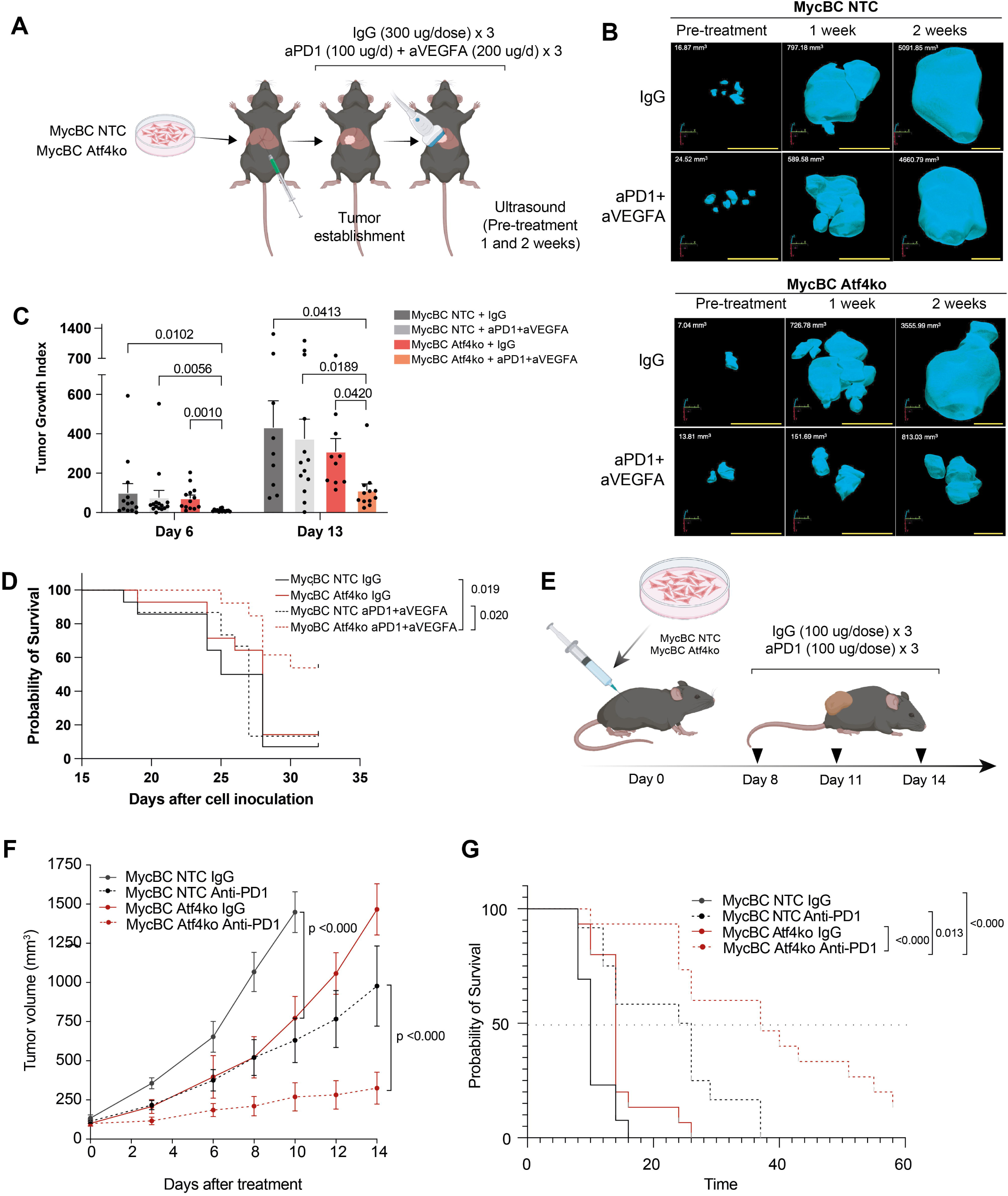
ATF4 deficiency enhances the efficacy of immunotherapy and prolongs survival in orthotopic and subcutaneous models of hepatocellular carcinoma. (A-B) Experimental scheme and three-dimensional ultrasound reconstructions of orthotopically implanted MycBC NTC and Atf4ko tumors (n = 15 mice/group). (C) Quantification of orthotopic tumor volume by ultrasound. Data are shown as mean ± SEM. Statistical significance was assessed using one-way ANOVA with Kruskal-Wallis test. (D) Survival analysis using Kaplan-Meier curves and Holm-Sidak multiple comparisons test. (E) Treatment protocol for subcutaneously inoculated MycBC NTC and Atf4Ko tumors (n = 12 mice/group). (F) Tumor growth curves with anti-PD1 treatment. Data are shown as mean ± SEM. Statistical significance was assessed using two-way ANOVA with multiple comparisons test and Uncorrected Fisher’s LSD. (G) Survival analysis using Kaplan-Meier curves and Holm-Sidak multiple comparisons test.

Given the marked reprogramming of the immune microenvironment observed in Atf4-deficient tumors, characterized by increased infiltration of effector T cells, NK cells, and antigen-presenting cells, alongside reduced macrophage and MDSC populations, we hypothesized that the loss of Atf4 may render tumors more susceptible to monotherapy with immune checkpoint blockade (ICB). To test this hypothesis, we treated C57Bl6J mice with subcutaneous tumors with anti-PD1 monotherapy (**Figure 6E**). While NTC tumors exhibited only a delay in tumor growth, similar to Atf4ko tumors without treatment, Atf4ko tumors showed a marked reduction in growth upon antiPD1 treatment (**Figure 6F**), indicating restored sensitivity to immune checkpoint inhibition. Importantly, tumor response translated into a significant increase in overall survival in mice bearing Atf4-deficient tumors (**Figure 6G**). These findings suggest that Atf4 contributes to an immunosuppressive tumor state that limits the efficacy of immune-checkpoint inhibition.

## Discussion

The suboptimal response rates (20-30%) of current immunotherapy combinations in HCC underscore the need for strategies that induce durable antitumor immunity (49–51). Stress response pathways, such as the UPR and ISR, are critical for tumor adaptation to metabolic and proteotoxic stress (52, 53). Our study provides the first systematic evaluation of the three UPR effectors -ATF4, ATF6, and XBP1- in HCC, establishing ATF4 as a primary driver of immune evasion in aggressive, immunotherapy-resistant models.

Using a model characterized by MYC overexpression and beta-catenin activating mutation - alterations present in ∼60% and ∼30% of HCC patients, respectively (54–57) -we identified a critical metabolic-immune axis. MYC is known to activate ATF4 via GCN2 to meet anabolic demands, with both factors co-occupying promoters of genes governing amino acid synthesis (37). Our findings expand this “cellular rheostat” model by showing that ATF4-null cells fail to upregulate rate-limiting enzymes like ALDH18A1 (P5CS), disrupting proline biosynthesis. Crucially, *Atf4* deletion reduced tumor growth only in immunocompetent mice, restoring sensitivity to immune checkpoint blockade. This indicates that the antitumor effect is mediated by a remodeled TME -characterized by increased effector T cells and a reduction in immunosuppressive macrophages- rather than intrinsic proliferation defects.

These findings are particularly relevant to *CTNNB1*-mutant HCCs, which are typically “cold” and resistant to anti-PD-1 therapy due to defective dendritic cell recruitment and impaired T cell activity (33). Since beta-catenin activation also promotes glutamine and proline metabolism (58, 59), ATF4 and beta-catenin appear to converge on the same biosynthetic programs. In MYC/beta-catenin-driven HCC, this creates a proline-rich, ammonia-reprogrammed microenvironment that fuels growth while sustaining immunosuppressive myeloid cells. *Atf4* deletion collapses these signals, converting “cold” tumors into “hot” ones amenable to immunotherapy.

While a recent study identified ATF4 as a driver of immune evasion in lung and pancreatic cancer via LCN2 (22), our HCC model suggests a distinct metabolic-centric mechanism. We did not observe significant Lcn2 expression, suggesting that ATF4-driven immunosuppression may be context-specific, with proline homeostasis representing a unique vulnerability in hepatocellular carcinoma.

Study limitations include the potential driver-specificity of the proline axis and the use of chronic CRISPR-mediated deletion. We observed that pharmacological ISR inhibition upstream (via DL233) does not phenocopy *Atf4* loss, likely because blocking eIF2alpha phosphorylation also triggers global translational re-activation. Nevertheless, ATF4 remains a uniquely attractive target; its inhibition simultaneously cripples tumor-intrinsic VEGFA-dependent angiogenesis and enhances the TME. Furthermore, attenuating ATF4 in T cells has been shown to reduce oxidative stress and promote TIL viability (60), positioning ATF4 inhibition as a powerful rational partner for immune checkpoint inhibitors.

## Abbreviations

ALDH18A1: Aldehyde Dehydrogenase 18 Family Member A1 (P5CS)
ALT: Alanine Aminotransferase
ASNS: Asparagine Synthetase
ATF4: Activating Transcription Factor 4
ATF6: Activating Transcription Factor 6
BMDM: Bone Marrow-Derived Macrophage
Casp3: Caspase 3
CCL5: C-C Motif Chemokine Ligand 5
CD4/CD8: Cluster of Differentiation 4/8
CHOP: C/EBP Homologous Protein (DDIT3)
CRISPR: Clustered Regularly Interspaced Short Palindromic Repeats
CTNNB1: Catenin Beta 1 (β-catenin)
DMEM: Dulbecco’s Modified Eagle Medium
ELISA: Enzyme-Linked Immunosorbent Assay
eIF2α: Eukaryotic Initiation Factor 2 Alpha
FACS: Fluorescence-Activated Cell Sorting
FBS: Fetal Bovine Serum
GCN2: General Control Nondenerepressible 2
GLUD1: Glutamate Dehydrogenase 1
GPT2: Glutamic-Pyruvic Transaminase 2
HCC: Hepatocellular Carcinoma
HGNC: HUGO Gene Nomenclature Committee
HPLC: High-Performance Liquid Chromatography
HRI: Heme-Regulated Inhibitor
ICI: Immune Checkpoint Inhibitor
IFNγ: Interferon Gamma
IL4: Interleukin 4
IL6: Interleukin 6
IRE1α: Inositol-Requiring Enzyme 1 Alpha
ISR: Integrated Stress Response
KO: Knockout
LCN2: Lipocalin 2
LPS: Lipopolysaccharide
MDSC: Myeloid-Derived Suppressor Cell
MTHFD2: Methylenetetrahydrofolate Dehydrogenase 2
MYC: Myelocytomatosis Oncogene
OAT: Ornithine Aminotransferase
P5CS: Delta-1-Pyrroline-5-Carboxylate Synthase
PD-1/PD-L1: Programmed Cell Death 1/Ligand 1
PERK: PKR-like Endoplasmic Reticulum Kinase
PKR: Protein Kinase RNA-activated
PRODH: Proline Dehydrogenase
PSPH: Phosphoserine Phosphatase
PYCR1: Pyrroline-5-Carboxylate Reductase 1
qRT-PCR: Quantitative Real-Time Polymerase Chain Reaction
sgRNA: Single Guide RNA
SHMT2: Serine Hydroxymethyltransferase 2
TCGA: The Cancer Genome Atlas
TIL: Tumor-Infiltrating Lymphocyte
TME: Tumor Microenvironment
TNFα: Tumor Necrosis Factor Alpha
uORF: Upstream Open Reading Frame
UPR: Unfolded Protein Response
VEGFA: Vascular Endothelial Growth Factor A
WT: Wild-Type
XBP1: X-box Binding Protein 1

## Financial support

SI was funded by a PhD scholarship by the School of Human Medicine of the University of Piura (Lima, Perú). JA’s work was supported by grants from the Instituto de Salud Carlos III (PI20/01663, PI25/00842 and PMP22/00054), the Asociación Española Contra el Cáncer (AECC; PRYCO234831REIG), and the Fundación Echebano.

## Conflict of interest

BS; reports grants from Bristol Myers Squibb and Roche; consulting fees from AstraZeneca, Bayer, Boston Scientific, Bristol Myers Squibb, Eisai, Incyte, Roche/Genentech, Sanofi Pasteur, and Sirtex Medical; honoraria from AstraZeneca, Eisai, Incyte, and Roche; and travel support from AstraZeneca, Bristol Myers Squibb, Eisai, Roche, and Sirtex Medical. JA has received fees for conferences and/or travel support from AstraZeneca, Pfizer, Bristol Myers Squibb and Roche. CEA has received fees for conferences and travel support from AstraZeneca and Roche.

No potential conflict of interest was reported by other authors.

## Authors’ contributions

JA established the hypothesis, designed the strategy and supervised the work with the help of JB and AU. JB, PS and PF, contributed to study concept and design. SI and ESM did most of the experimental work with AF, SA, NH, AU, AM, GGP, EM, CA, DLL, completing specific experiments and helping with the acquisition of data. Analysis and interpretation of experimental and statistical data was performed by SI, ESM, AU, JB, PS, PF and JA. GGP, CEA and EM performed the immunohistochemistry of HSPA5 and the multiplex immunofluorescences in animals. The bioinformatic analyses were performed by JB, AF and JA with the help of SB and AM. SI, AU, JB and JA drafted the manuscript. PS, PF, BS and YF critically revised the manuscript. The manuscript was supported with funding obtained from JA and BS.

## Data availability statement

The gsadapt pipeline is available at https://github.com/unav-hcclab/gsadapt.git (23). RNA sequencing data of human HCC in the form of expression matrices were downloaded from the Integrative Molecular Database of Hepatocellular Carcinoma (HCCDB) with references HCCDB25, HCCDB15, and HCCDB18, for OEP000321 (OEP), The Cancer Genome Atlas Liver Hepatocellular Carcinoma (TCGA-LIHC, TCGA), and the International Cancer Genome Consortium Japan Liver Cancer-RIKEN (ICGC-LIRI-JP, LIRI), respectively. and can be found here: http://www.lifeome.net/database/hccdb/home.html. Single-cell RNA sequencing (scRNA-seq) data were obtained from the Gene Expression Omnibus (GEO) under accession GSE149614 (29). All custom scripts and RNA sequencing data from experimental models are available upon reasonable request.

## Acknowledgements

We thank to Diego Alignani and the Flow Cytometry Platform at CIMALABS Diagnostics. We thank Rubió metabolomics for their advice on data analysis. We thank the help of the Genomics Platform (Francesco Marchese, Jon Zazpe and Annarosaria De Vito), Morphology and Imaging Platform (Laura Guembe, Xabier Morales) of CIMA University of Navarra, for their unconditional support. We thank the Biobank of the University of Navarra (Virginia Villar, Antonia Fortuño).

## References

1. Bray F, Laversanne M, Sung H, Ferlay J, Siegel RL, Soerjomataram I, et al. Global cancer statistics 2022: GLOBOCAN estimates of incidence and mortality worldwide for 36 cancers in 185 countries. CA Cancer J Clin. 2024;74(3):229–63.

2. Ladd AD, Duarte S, Sahin I, Zarrinpar A. Mechanisms of drug resistance in HCC. Hepatology. 2024;79(4):926–40.

3. Kou L, Xie X, Chen X, Li B, Li J, Li Y. The progress of research on immune checkpoint inhibitor resistance and reversal strategies for hepatocellular carcinoma. Cancer Immunol Immunother. 2023;72(12):3953–69.

4. Wang Z, Wang Y, Gao P, Ding J. Immune checkpoint inhibitor resistance in hepatocellular carcinoma. Cancer Lett. 2023;555:216038.

5. Kim LC, Lesner NP, Simon MC. Cancer Metabolism under Limiting Oxygen Conditions. Cold Spring Harb Perspect Med. 2024;14(2).

6. Kim SJ, Khadka D, Seo JH. Interplay between Solid Tumors and Tumor Microenvironment. Front Immunol. 2022;13:882718.

7. Paredes F, Williams HC, San Martin A. Metabolic adaptation in hypoxia and cancer. Cancer Lett. 2021;502:133–42.

8. Tameire F, Verginadis, II, Koumenis C. Cell intrinsic and extrinsic activators of the unfolded protein response in cancer: Mechanisms and targets for therapy. Semin Cancer Biol. 2015;33:3–15.

9. Oakes SA. Endoplasmic Reticulum Stress Signaling in Cancer Cells. Am J Pathol. 2020;190(5):934–46.

10. Obacz J, Avril T, Rubio-Patino C, Bossowski JP, Igbaria A, Ricci JE, et al. Regulation of tumor-stroma interactions by the unfolded protein response. FEBS J. 2019;286(2):279–96.

11. Pakos-Zebrucka K, Koryga I, Mnich K, Ljujic M, Samali A, Gorman AM. The integrated stress response. EMBO Rep. 2016;17(10):1374–95.

12. Pathak SS, Liu D, Li T, de Zavalia N, Zhu L, Li J, et al. The eIF2alpha Kinase GCN2 Modulates Period and Rhythmicity of the Circadian Clock by Translational Control of Atf4. Neuron. 2019;104(4):724–35 e6.

13. Wortel IMNvdMLTKMSvLFN. Surviving Stress: Modulation of ATF4-Mediated Stress Responses in Normal and Malignant Cells. Trends in Endocrinology and Metabolism. 2017;28(11):794–806.

14. He F, Zhang P, Liu J, Wang R, Kaufman RJ, Yaden BC, et al. ATF4 suppresses hepatocarcinogenesis by inducing SLC7A11 (xCT) to block stress-related ferroptosis. J Hepatol. 2023;79(2):362–77.

15. DeZwaan-McCabe D, Riordan JD, Arensdorf AM, Icardi MS, Dupuy AJ, Rutkowski DT. The stress-regulated transcription factor CHOP promotes hepatic inflammatory gene expression, fibrosis, and oncogenesis. PLoS Genet. 2013;9(12):e1003937.

16. Zheng Y, Zhou Q, Zhao C, Li J, Yu Z, Zhu Q. ATP citrate lyase inhibitor triggers endoplasmic reticulum stress to induce hepatocellular carcinoma cell apoptosis via p-eIF2alpha/ATF4/CHOP axis. J Cell Mol Med. 2021;25(3):1468–79.

17. Li R, Wang K, Hou N, Tian Y, Gong B, Tang M. Activating transcription factor 4-mediated upregulation of Heat Shock Protein Family A Member 4 promotes hepatocellular carcinoma progression through activation of Wnt/beta-catenin/EMT signaling pathway. Int J Biol Macromol. 2025;333(Pt 2):148850.

18. Zhang Z, Yin J, Zhang C, Liang N, Bai N, Chang A, et al. Activating transcription factor 4 increases chemotherapeutics resistance of human hepatocellular carcinoma. Cancer Biol Ther. 2012;13(6):435–42.

19. Dai Z, Wang X, Peng R, Zhang B, Han Q, Lin J, et al. Induction of IL-6Ralpha by ATF3 enhances IL-6 mediated sorafenib and regorafenib resistance in hepatocellular carcinoma. Cancer Lett. 2022;524:161–71.

20. Hetz C, Zhang K, Kaufman RJ. Mechanisms, regulation and functions of the unfolded protein response. Nat Rev Mol Cell Biol. 2020;21(8):421–38.

21. Chevet E, Hetz C, Samali A. Endoplasmic reticulum stress-activated cell reprogramming in oncogenesis. Cancer Discov. 2015;5(6):586–97.

22. Bossowski JP, Pillai R, Kilian J, Wong Lau A, Nakamura M, Rashidfarrokhi A, et al. The integrated stress response promotes immune evasion through lipocalin 2. Nature. 2026.

23. Barace S, Santamaria E, Infante S, Arcelus S, De La Fuente J, Goni E, et al. Application of Graph Models to the Identification of Transcriptomic Oncometabolic Pathways in Human Hepatocellular Carcinoma. Biomolecules. 2024;14(6).

24. Villanueva A. Hepatocellular Carcinoma. N Engl J Med. 2019;380(15):1450–62.

25. Pfister D, Nunez NG, Pinyol R, Govaere O, Pinter M, Szydlowska M, et al. NASH limits anti-tumour surveillance in immunotherapy-treated HCC. Nature. 2021;592(7854):450–6.

26. Henkel AS. Unfolded Protein Response Sensors in Hepatic Lipid Metabolism and Nonalcoholic Fatty Liver Disease. Semin Liver Dis. 2018;38(4):320–32.

27. Lazar C, Uta M, Branza-Nichita N. Modulation of the unfolded protein response by the human hepatitis B virus. Front Microbiol. 2014;5:433.

28. Malhi H, Kaufman RJ. Endoplasmic reticulum stress in liver disease. J Hepatol. 2011;54(4):795–809.

29. Lu Y, Yang A, Quan C, Pan Y, Zhang H, Li Y, et al. A single-cell atlas of the multicellular ecosystem of primary and metastatic hepatocellular carcinoma. Nat Commun. 2022;13(1):4594.

30. Iwakoshi NN, Lee AH, Vallabhajosyula P, Otipoby KL, Rajewsky K, Glimcher LH. Plasma cell differentiation and the unfolded protein response intersect at the transcription factor XBP-1. Nat Immunol. 2003;4(4):321–9.

31. Hoshida Y, Villanueva A, Kobayashi M, Peix J, Chiang DY, Camargo A, et al. Gene expression in fixed tissues and outcome in hepatocellular carcinoma. N Engl J Med. 2008;359(19):1995–2004.

32. Hoshida Y, Nijman SM, Kobayashi M, Chan JA, Brunet JP, Chiang DY, et al. Integrative transcriptome analysis reveals common molecular subclasses of human hepatocellular carcinoma. Cancer Res. 2009;69(18):7385–92.

33. Ruiz de Galarreta M, Bresnahan E, Molina-Sanchez P, Lindblad KE, Maier B, Sia D, et al. beta-Catenin Activation Promotes Immune Escape and Resistance to Anti-PD-1 Therapy in Hepatocellular Carcinoma. Cancer Discov. 2019;9(8):1124–41.

34. Hetz C, Papa FR. The Unfolded Protein Response and Cell Fate Control. Mol Cell. 2018;69(2):169–81.

35. Lin JH, Li H, Yasumura D, Cohen HR, Zhang C, Panning B, et al. IRE1 signaling affects cell fate during the unfolded protein response. Science. 2007;318(5852):944–9.

36. Argemi J, Kress TR, Chang HCY, Ferrero R, Bertolo C, Moreno H, et al. X-box Binding Protein 1 Regulates Unfolded Protein, Acute-Phase, and DNA Damage Responses During Regeneration of Mouse Liver. Gastroenterology. 2017;152(5):1203–16 e15.

37. Tameire F, Verginadis, II, Leli NM, Polte C, Conn CS, Ojha R, et al. ATF4 couples MYC-dependent translational activity to bioenergetic demands during tumour progression. Nat Cell Biol. 2019;21(7):889–99.

38. Liu B, Chen P, Xi D, Zhu H, Gao Y. ATF4 regulates CCL2 expression to promote endometrial cancer growth by controlling macrophage infiltration. Exp Cell Res. 2017;360(2):105–12.

39. Wang Y, Alam GN, Ning Y, Visioli F, Dong Z, Nor JE, et al. The unfolded protein response induces the angiogenic switch in human tumor cells through the PERK/ATF4 pathway. Cancer Res. 2012;72(20):5396–406.

40. Liu W, Hancock CN, Fischer JW, Harman M, Phang JM. Proline biosynthesis augments tumor cell growth and aerobic glycolysis: involvement of pyridine nucleotides. Sci Rep. 2015;5:17206.

41. Ding Z, Ericksen RE, Escande-Beillard N, Lee QY, Loh A, Denil S, et al. Metabolic pathway analyses identify proline biosynthesis pathway as a promoter of liver tumorigenesis. J Hepatol. 2020;72(4):725–35.

42. Su P, Yang Y, Zheng H. The “serine code” of metabolic reprogramming: multidimensional roles of the serine synthesis pathway in tumors and novel breakthroughs for targeted therapy. Front Immunol. 2026;17:1779543.

43. Wei W, Li H, Tian S, Zhang C, Liu J, Tao W, et al. Asparagine drives immune evasion in bladder cancer via RIG-I stability and type I IFN signaling. J Clin Invest. 2025;135(8).

44. Kimura T, Nada S, Takegahara N, Okuno T, Nojima S, Kang S, et al. Polarization of M2 macrophages requires Lamtor1 that integrates cytokine and amino-acid signals. Nat Commun. 2016;7:13130.

45. Chang L, Zhou L, Jiang S, Wei K, Li J, Zhao J, et al. Proline-Mediated Inhibition of ATPIF1-mTOR Signaling Alleviates Radiation-Induced Macrophage Polarization and Colon Inflammation. Inflammation. 2026;49(1).

46. Liu Y, Xun Z, Ma K, Liang S, Li X, Zhou S, et al. Identification of a tumour immune barrier in the HCC microenvironment that determines the efficacy of immunotherapy. J Hepatol. 2023;78(4):770–82.

47. Wei CY, Zhu MX, Zhang PF, Huang XY, Wan JK, Yao XZ, et al. PKCalpha/ZFP64/CSF1 axis resets the tumor microenvironment and fuels anti-PD1 resistance in hepatocellular carcinoma. J Hepatol. 2022;77(1):163–76.

48. Zhang X, Lao M, Sun K, Yang H, He L, Liu X, et al. Sphingolipid synthesis in tumor-associated macrophages confers immunotherapy resistance in hepatocellular carcinoma. Sci Adv. 2025;11(21):eadv0558.

49. Finn RS, Qin S, Ikeda M, Galle PR, Ducreux M, Kim T-Y, et al. IMbrave150: Updated overall survival (OS) data from a global, randomized, open-label phase III study of atezolizumab (atezo) + bevacizumab (bev) versus sorafenib (sor) in patients (pts) with unresectable hepatocellular carcinoma (HCC). Journal of Clinical Oncology. 2021;39(3_suppl):267-.

50. Sangro B, Chan SL, Kelley RK, Lau G, Kudo M, Sukeepaisarnjaroen W, et al. Four-year overall survival update from the phase III HIMALAYA study of tremelimumab plus durvalumab in unresectable hepatocellular carcinoma. Ann Oncol. 2024;35(5):448–57.

51. Yau T, Galle PR, Decaens T, Sangro B, Qin S, da Fonseca LG, et al. Nivolumab plus ipilimumab versus lenvatinib or sorafenib as first-line treatment for unresectable hepatocellular carcinoma (CheckMate 9DW): an open-label, randomised, phase 3 trial. Lancet. 2025;405(10492):1851–64.

52. Chen X, Cubillos-Ruiz JR. Endoplasmic reticulum stress signals in the tumour and its microenvironment. Nat Rev Cancer. 2021;21(2):71–88.

53. Cubillos-Ruiz JR, Bettigole SE, Glimcher LH. Tumorigenic and Immunosuppressive Effects of Endoplasmic Reticulum Stress in Cancer. Cell. 2017;168(4):692–706.

54. Sanchez-Vega F, Mina M, Armenia J, Chatila WK, Luna A, La KC, et al. Oncogenic Signaling Pathways in The Cancer Genome Atlas. Cell. 2018;173(2):321–37 e10.

55. Schulze K, Imbeaud S, Letouze E, Alexandrov LB, Calderaro J, Rebouissou S, et al. Exome sequencing of hepatocellular carcinomas identifies new mutational signatures and potential therapeutic targets. Nat Genet. 2015;47(5):505–11.

56. Guichard C, Amaddeo G, Imbeaud S, Ladeiro Y, Pelletier L, Maad IB, et al. Integrated analysis of somatic mutations and focal copy-number changes identifies key genes and pathways in hepatocellular carcinoma. Nat Genet. 2012;44(6):694–8.

57. Chen L, Zhang C, Xue R, Liu M, Bai J, Bao J, et al. Deep whole-genome analysis of 494 hepatocellular carcinomas. Nature. 2024;627(8004):586–93.

58. Ren X, Rong Z, Liu X, Gao J, Xu X, Zi Y, et al. The Protein Kinase Activity of NME7 Activates Wnt/beta-Catenin Signaling to Promote One-Carbon Metabolism in Hepatocellular Carcinoma. Cancer Res. 2022;82(1):60–74.

59. Wang Y, Cheng C, Lu Y, Lian Z, Liu Q, Xu Y, et al. beta-Catenin Activation Reprograms Ammonia Metabolism to Promote Senescence Resistance in Hepatocellular Carcinoma. Cancer Res. 2024;84(10):1643–58.

60. Alicea Pauneto CDM, Riesenberg BP, Gandy EJ, Kennedy AS, Clutton GT, Hem JW, et al. Intra-tumoral hypoxia promotes CD8(+) T cell dysfunction via chronic activation of integrated stress response transcription factor ATF4. Immunity. 2025;58(10):2489–504 e8.

