## Supplementary Information for "ATF4 programs proline-dependent immune evasion in β-Catenin-driven hepatocellular carcinoma"

**Supplementary Materials and Methods**

**Single-sample gene set enrichment analysis (ssGSEA)**

ssGSEA was performed using the *corto* R package (v1.2.4) on log^2^(CPM+1)-transformed expression matrices. Enrichment scores were z-score normalized across samples within each cohort and compared between tumor and non-tumor groups using the Wilcoxon rank-sum test with Benjamini-Hochberg (BH) correction. Analyses used *tidyverse* (v2.0.0), *rstatix* (v0.7.2), and *ggplot2* (v3.4.4) in R v4.3.

**Single-cell RNA-seq dataset acquisition and bioinformatic analysis**

Publicly available single-cell RNA sequencing (scRNA-seq) data were obtained from the Gene Expression Omnibus (GEO) under accession GSE149614, corresponding to a previously published study (1) study. This dataset comprises >70,000 single-cell transcriptomes generated from 10 patients with hepatocellular carcinoma (HCC), including samples from primary tumor (PT), portal vein tumor thrombus (PVTT), metastatic lymph node (MLN), and matched non-tumor liver (NTL). Libraries were prepared using the 10x Genomics Chromium Single Cell 3′ v2 chemistry and sequenced on an Illumina NovaSeq 6000 platform. Raw sequencing reads were originally processed by the study authors using Cell Ranger v2.2.0 with alignment to the human reference genome hg38. For the present analysis, the processed count matrix and accompanying metadata were imported into Seurat (R environment). A sparse gene-by-cell expression matrix was used to construct a Seurat object. Quality-control filtering excluded cells with fewer than 200 detected genes, more than 6,000 detected genes, or >10% mitochondrial transcripts. To optimize computational efficiency while preserving cellular diversity, a random subset of up to 20,000 cells was retained for downstream analyses. Data were log-normalized, highly variable genes were identified (n = 3,000), and scaled expression values were subjected to principal component analysis (PCA). Uniform Manifold Approximation and Projection (UMAP) was performed using the first 20 principal components, followed by shared nearest-neighbor graph construction and unsupervised clustering (resolution = 0.5). Cell identities were assigned according to the provided annotations. Signature enrichment was calculated using AUCell, which ranks genes within each cell and computes the area under the recovery curve (AUC) for genes present in each signature. AUC scores were added to cell metadata and compared across cell populations. Visualization outputs included UMAP embeddings, bubble plots summarizing mean z-scored AUCell activity and fraction of enriched cells. Multi-panel figures were generated in R using ggplot2 and patchwork.

**Survival analysis and clinical variable association**

For survival analysis, tumor samples were stratified by ssGSEA score quartiles per signature, and patients in Q1 (lowest) and Q4 (highest) were compared. Overall survival was estimated by Kaplan-Meier and differences assessed by log-rank test. Hazard ratios with 95% CIs were derived from univariate Cox regression (*survival*, v3.5) and displayed as forest plots and Kaplan-Meier curves (*survminer*, v0.4.9). Associations between Q1/Q4 stratification and categorical clinical variables (BCLC stage, TNM stage, AFP, etiology, vascular invasion, *CTNNB1*/*TP53* status, fibrosis) were assessed by Fisher’s exact test or chi-squared test, as appropriate.

**Pseudotime trajectory inference**

Pseudotime trajectory inference was performed using the Slingshot algorithm (*slingshot*, v2.8.0) on ssGSEA enrichment scores for all refined signatures. Tumor and non-tumor samples were combined, with non-tumor samples set as the trajectory origin. Each sample was assigned a pseudotime value along the inferred trajectory, and tumor samples were classified as “early” or “late” based on the median tumor pseudotime. Distributions were visualized as histograms and violin plots. Group comparisons were performed using the Wilcoxon rank-sum test, and early vs late classification was assessed for prognostic relevance by Kaplan-Meier and Cox regression, following the same clinical variable association approach as above.

**Differential expression and gene set enrichment analysis**

Differential gene expression between early and late pseudotime groups, and between late tumors and non-tumor tissue, was performed per cohort using *limma* (v3.56). Genes with CPM >1 in ≥20% of samples were retained, and linear models were fitted with *lmFit()* followed by *eBayes()* moderation. Differentially expressed genes were defined by |log^2^FC| >0.585 and BH-adjusted p<0.05. Cross-cohort overlap was visualised using Venn diagrams (*VennDiagram*, v1.7.3) and UpSet plots (*UpSetR*, v1.4.0).

Pre-ranked GSEA was performed on the early vs late comparison using *fgsea* (v1.26.0, 10,000 permutations), with genes ranked by log^2^FC. Gene sets from MSigDB (v2023.2) were retrieved via *msigdbr*, including Hallmark (H), curated (C2), oncogenic (C6), and immunologic (C7) collections. Pathways with FDR-adjusted p<0.05 were considered significant.

**Immune cell deconvolution**

Immune cell composition was estimated by computational deconvolution using the MIXTURE algorithm, applied independently to each cohort. Resulting cell type proportions were z-score normalized within each cohort. Pearson correlations between ssGSEA scores and cell type proportions, and between pseudotime values and cell type proportions, were computed using *corr.test()* from *psych* (v2.3.9). Differences between early and late groups were assessed by Wilcoxon rank-sum test and visualized as heatmaps and boxplots using *pheatmap* (v1.0.12) and *ggpubr* (v0.6.0).

**Individual gene expression analysis**

Expression of individual UPR/ISR genes (*ATF4*, *ATF6*, *XBP1*, *ERN1*, *EIF2AK1–EIF2AK4*) was compared between tumor and non-tumor tissue across cohorts. Log^2^ fold changes were calculated relative to the non-tumor mean and differences assessed by Wilcoxon rank-sum test with FDR correction, visualized as box-and-violin plots (*ggplot2*, *patchwork* v1.1.3).

**Immunohistochemistry of human liver samples**

Formalin-fixed, paraffin-embedded (FFPE) HCC sections (from ALD and MAFLD-related HCC) were stained for BiP/HSPA5 using the Ventana BenchMark automated platform (Roche). Antigen retrieval was performed with CC1 buffer (Tris-EDTA, pH 8.2) for 36 min at 95°C. Endogenous peroxidase was blocked using Ultraview DAB Inhibitor (4 min, 37°C). Sections were incubated with primary anti-BiP/HSPA5 antibody (Abcam ab108613, 1:100) for 32 min. Signal was detected using the Ultraview Universal DAB Detection Kit according to standard protocols. Slides were counterstained with hematoxylin and reviewed alongside H&E-stained sections.

**Cell lines, culture and vectors**

The HCC PM299L cell line provided Dr A. Lujambio (MSSM, NY, USA), has been previously described (2). It was obtained from a tumor generated in mice by hydrodynamic injection of plasmids MYC-lucOS (encoding MYC, luciferase and OVA peptides 257–264 and 323–339) and CTNNB1 (encoding a constitutive active beta-catenin form). Along the manuscript the cell line has been named MycBC. These cells were cultured in complete medium: DMEM medium high glucose + pyruvate (GIBCO-41966-029) containing 5% heat-inactivated fetal bovine serum (FBS),1% penicillin/streptomycin (GIBCO -15140122) and 0.1% β-mercaptoetanol (GIBCO-A331350010). For amino acid deprivation stress conditions, cell lines were cultured in DMEM medium high glucose (-L-glutamine, -pyruvate, -L-methionine, -L-cystine) (GIBCO-21013-024), containing 1% FBS, 1% penicillin-streptomycin and 0.1% β-mercaptoetanol during 8 hours. For inhibition of N-glycosylation stress conditions, cell lines were cultured in DMEM complete medium and were treated with Tunicamycin (MP Biomedicals-215002810) 1 ug/mL during 8 hours. All the cell lines were maintained at 37ºC and 5% CO2.

**Cell proliferation assay**

Cell proliferation was assessed using the 3-(4,5-dimethylthiazol-2-yl)-5-(3-carboxymethoxyphenyl)-2-(4-sulfophenyl)-2H-tetrazolium (MTS) colorimetric assay according to the manufacturer's instructions (CellTiter 96® AQueous One Solution, Promega). Cells were seeded in 96-well plates at a density of 20,000 cells per well in 100 µL of DMEM complete medium and incubated for 24 h at 37 °C in a humidified atmosphere with 5% CO₂. Subsequently, the cells were subjected to the corresponding treatment for 6,14 and 24 hours. After treatment, the culture medium was renewed, and 20 µL of the MTS reagent were added directly to each well. The plates were incubated for 1–3 h at 37 °C. Finally, quantification was performed by measuring the absorbance at 490 nm using a microplate reader (GloMax® Discover-Promega).

**Apoptosis**

HCC tumor cell lines (WT and UPR TF KO) were seeded at a density of 1 million cells per 6-well. After 24 h, media was changed and cells were cultured under basal or stress conditions (tunicamycin or aminoacid deprivation) for 8 h. Finally, cells were harvested and analysed for apoptosis, by staining with Propidium iodide and annexin V (BD Pharmingen™ FITC Annexin V, 556419). The manufacturer's instructions were followed, and all assays were evaluated by flow cytometry.

**Cell cycle analysis**

HCC tumor cell lines (WT and UPR TF KO) were seeded at a density of 1 million cells per 6-well. After 24 h, media was changed and cells were cultured under basal or stress conditions (tunicamycin or aminoacid deprivation) for 8 h. Cells were harvested and stained with a propidium iodide solution (10 mL 1X PBS, 20 µL of 100 mg/mL RNase A, and 200 µL of 1 mg/mL PI) to evaluate the cell cycle. All samples were analysed by flow cytometry.

**RNA sequencing of cell lines**

RNA was subjected to quantity and quality control using Qubit HS RNA Assay Kit (Thermo Fisher Scientific) and 4200 Tapestation with High Sensitivity RNA ScreenTape (Agilent Technologies). All RNA samples were high-quality, with RIN values higher than 8. Library preparation was performed using the Illumina Stranded mRNA Prep Ligation kit (Illumina) following the manufacturer’s protocol. All sequencing libraries were constructed from 100 ng of total RNA according to the manufacturer's instruction. Briefly, the protocol selects and purifies poly(A) containing RNA molecules using magnetic beads coated with poly(T) oligos. Poly(A)-RNAs are fragmented and reverse transcribed into first cDNA strand using random primers. The second cDNA strand is synthesized in the presence of dUTP to ensure strand specificity. Resulting cDNA fragments are purified with AMPure XP beads (Beckman Coulter), adenylated at 3′ ends and then ligated with uniquely indexed sequencing adapters. Ligated fragments are purified and PCR amplified to obtain the final libraries. The quality and quantity of the libraries were verified using Qubit dsDNA HS Assay Kit (Thermo Fisher Scientific) and 4200 Tapestation with High Sensitivity D1000 ScreenTape (Agilent Technologies). Libraries were then sequenced using a NextSeq2000 sequencer (Illumina). 20-30 million pair-end reads (50 bp x2) were sequenced for each sample and demultiplexed using bcl2fastq.

**Imaging of orthotopic tumors**

Tumor progression in mice was longitudinally evaluated by high-frequency transabdominal ultrasonography at days 11 (pre-treatment, for randomization), 17 and 24 post-implantation using the Vevo3100 imaging platform (VisualSonics, Toronto, Canada) equipped with an MX550 linear-array transducer (central frequency 40 MHz; focal depth 7.0 mm). B-mode acquisitions were performed at a frame rate of 557 frames per second, providing a maximum two-dimensional (2D) FoV of 14.1 × 15.0 mm. The system afforded a spatial resolution of approximately 90 μm laterally and 40 μm axially, enabling high-resolution visualization of tumor architecture and boundary definition suitable for volumetric reconstruction.

Animals were anesthetized with 2% isoflurane (Isoflo®, ABBOTT, Spain) delivered in 80% oxygen and positioned prone on a temperature-controlled heating platform to maintain physiological homeostasis. Electrocardiogram (ECG) electrodes were placed on the limbs for continuous monitoring of cardiac rhythm, and respiratory rate was simultaneously recorded to ensure stable anesthetic depth and minimize motion artifacts. Abdominal hair was removed using a depilatory cream (Veet, Reckitt, France), and pre-warmed acoustic coupling gel (Quick Eco-Gel, Lessa, Spain) was applied prior to imaging.

Following tumor localization in B-mode, three-dimensional (3D) datasets were automatically acquired using a motorized linear translation stage integrated into the imaging system. Cardiorespiratory signals were synchronized with image acquisition to reduce motion-related distortions and improve volumetric accuracy. Tumor volumes (mm³) were quantified offline using VevoLab software (version 5.8.1, VisualSonics, Canada). A trained operator performed semi-automated segmentation by delineating a region of interest (ROI) along the tumor margins across sequential image slices, enabling reconstruction of the tumor volume through stacked-planar integration.

**Amino acid profiling**

HCC UPR TF Ko tumor cells were seeded at a density of 6 x 10⁶ in 100 mm plates in DMEM medium high glucose + pyruvate containing 5% dialyzed fetal bovine serum (dFBS), 1% penicillin/streptomycin and 0.1% β-mercaptoetanol. After 24 h of culture, the seeding medium was renewed, and after 8 h the supernatants were collected. Metabolic amino acid profiling of the supernatants was carried out by [Metabolomics](https://www.sciencedirect.com/topics/biochemistry-genetics-and-molecular-biology/metabolomics) S.L. Rubió. Multiple ultra-high performance liquid chromatography coupled to mass spectrometry (UHPLC-MS) platforms were employed to quantify analytes in the samples.

**Multiplex immunofluorescence**

Tumor samples were processed for histological analysis by preparing 4 µm thick sections mounted on slides. Immune cell infiltration was assessed by multiplex immunofluorescence staining with a lymphoid panel including CD4, CD8, Foxp3, CD20, Ki67, and Granzyme B. In parallel, a myeloid panel composed of CD11c, CD11b, Ly6G, and F4/80 was used to characterize myeloid cell populations in the tumor microenvironment. The data were analyzed by QuPath (Bankhead, P. et al. QuPath: Open source software for digital pathology image analysis. Scientific Reports (2017). <https://doi.org/10.1038/s41598-017-17204-5>) for cell visualization and phenotyping and CellPose (Stringer, C., Wang, T., Michaelos, M., & Pachitariu, M. (2021). Cellpose: a generalist algorithm for cellular segmentation. Nature methods, 18(1), 100-106.) for cell segmentation

**Flow Cytometry**

Tumor samples were enzymatically dissociated using collagenase and DNase (Roche) for 15 minutes, followed by mechanical homogenization. The resulting cells were incubated with Fc Block™ (Biolegend) for 15 minutes. Cells were subsequently stained with the indicated antibody panels (see Supplementary Methods) to identify lymphoid and myeloid populations. Non-viable cells were excluded using Promofluor Invitrogen™ LIVE/DEAD™ Fixable Near IR-876 (Live/Dead staining). Data acquisition was performed on a CytoFLEX LX flow cytometer (Beckman Coulter), and analyses were conducted using FlowJo software (v10.10.0). The gating strategy is provided in the Supplementary Figure 8 and the antibodies used are in Supplementary Table S4. For cell staining (tumor cell lines and macrophages), they are washed with Phosphate-Buffered Saline (PBS) 1X and collected with a scraper, the pellet is resuspended in 200 uL of PBS 1X and the cytometry staining steps are followed.

**Software and data availability**

All analyses were performed in R (v4.3.3). The *gsadapt* pipeline is available at <https://github.com/unav-hcclab/gsadapt.git>. All custom scripts are available upon reasonable request.

**Supplementary Tables**

**Supplementary Table 1: sgRNA sequences used in CRISPR/Cas9**

| **UPR Transcription factor** | **System** | **Sequence** | **knocked out** |
| --- | --- | --- | --- |
| moXBP1 | sgRNA 1 | GCTCACGCACCTGAGCCCGG | Selected |
|  | sgRNA 2 | GGACACGCTGGATCCTGACG | No selected |
|  | sgRNA 3 | TCTGGCCAGCCCGCCTCCGG | No selected |
| moATF4 | sgRNA 1 | CAAACCCGACTGGTCGAAGG | No selected |
|  | sgRNA 2 | TTGTCGCTGGAGAACCCATG | Selected |
|  | sgRNA 3 | TCTCTTAGATGACTATCTGG | No selected |
| moATF6 | sgRNA 1 | CATGGACCAGATGAAGACTG | No selected |
|  | sgRNA 2 | GAGTCAGACCTATGGAGCCC | Selected |
|  | sgRNA 3 | GAACACGAGTCTGTGGACCG | No selected |

**Supplementary Table 2: Oligonuclotide sequences**

| **Gene Symbol** | **Sense** | **Sequence** |
| --- | --- | --- |
| Edem1 | Foward | GTGCCCAGATGCGCGACCTG |
|  | Reverse | CTATGAGCAGAAAGGAGGCTTCCC |
| Dnabj9 | Foward | TCGTCTTTGCAATCTGCATT |
|  | Reverse | GGCATCCGAGAGTGTTTCAT |
| Dnabj11 | Foward | GCATGGAGTACCCCTTTATTGGAGAAGG |
|  | Reverse | TTCCACAGCTTGGCTCCTGGCC |
| Angptl6 | Foward | AGACTCCCTCTCTTGGCACA |
|  | Reverse | CAGTAGACCCCGTCCTGGTA |
| Trib3 | Foward | TTTGTCTTCAGCAACTGTGAGAGGACGAA |
|  | Reverse | CCTCAGGCAGGGCAAAGGTCC |
| Ddit3 (Chop) | Foward | GCTGGGAGCTGGAAGCCTGGTATG |
|  | Reverse | TCCCTGGTCAGGCGCTCGATTTCC |
| Bip | Foward | TCGTCTTTGCAATCTGCATT |
|  | Reverse | GGCATCCGAGAGTGTTTCAT |
| Grp94 | Foward | CAGGACAAAATCTACTTCATGGCTGGGTC |
|  | Reverse | ACGGGCAAGGACATCTCTACAAATTACTATGC |

**Supplementary Table 3: Antibodies used for Western Blot**

| **Antibody** | **Supplier** | **Reference** | **Dilution** | **Band Size (Kda)** |
| --- | --- | --- | --- | --- |
| ATF4 | CELL SIGNALLING | 11815S | 1:1000 | 50 |
| XBP1S | CELL SIGNALLING | 12782S | 1:1000 | 50 |
| ATF6 | PROTEINTECH | 24169-1-AP | 1:1000 | 60-90 |
| Pperk | CELL SIGNALLING | 3179s | 1:1000 | 150 |
| Perk | CELL SIGNALLING | 5683s | 1:1000 | 150 |
| P-EIF2a | CELL SIGNALLING | 9721s | 1:1000 | 37 |
| VINCULINA | CELL SIGNALLING | 13901S | 1:1001 | 126 |
| SQSTM1/p62 | CELL SIGNALLING | 39749S | 1:1000 | 62 |
| PUROMICINA 12D10 | SIGMA ALDRICH | MABE343 | 1:500 |  |
| pS6 - serine235 | CELL SIGNALLING | 4857 | 1:1000 | 32 |
| LC3B D11 | CELL SIGNALLING | 3868 | 1:1000 | 15-20 |

**Supplementary Table 4: Antibodies used for flow cytometry**

| **Targeted Molecule** | **Fluorochrome** | **Working Dilution (1:X)** | **Clone** | **Supplier** | **Reference** |
| --- | --- | --- | --- | --- | --- |
| Fc Block | Isotype Rat IgG2a | 100 | 93 | BioLegend | 553142 |
| F4/80 | BV421 | 100 | BM8 | BioLegend | 123137 |
| CD11b | BUV661 | 500 | M1/70 | Invitrogen | 376-0112-82 |
| CD86 | BV785 | 600 | GL-1 | Biolegend | 105043 |
| MHC II | BUV395 | 100 | 2G9 | BD Biosciences | 569 244 |
| CD38 | PerCP/Cy5.5 | 400 | 90 | Biolegend | 102722 |
| Arg1 | PE | 400 | W21047I | Biolegend | 54165804 |
| CD206 | PE/Cy7 | 400 | C068C2 | Biolegend | 141720 |
| CD45 | APC-Cy7 | 200 | 30-F11 | BioLegend | 103116 |
| I-A^b^ | BUV 395 | 200 | 2G9 | BD Biosciences | 569 244 |
| CD11c | BV510 | 200 | N418 | Biolegend | 117353 |
| PD-L1 | RY610 | 100 | MIH5 | BD Biosciences | 758309 |
| Ly6C | (Alexa fluor 700) | 1000 |  |  |  |
| CD3e | BUV495 | 50 | 53-6.7 | BD Biosciences | 553031 |
| CD4 | Alexa700 | 1000 | RM4-5 | BioLegend | 100536 |
| CD8a | FITC | 200 | 53-6.7 | BD Biosciences | 553031 |
| CD25 | PE/Cy7 | 200 | 3C7 | Biolegend | 101916 |
| FoxP3 | BV421 | 200 | 150D | Biolegend | 320011 |
| Tet OVA | APC | 100 | - | MBL | TB-5001-2C |
| PD1 | PerCP/Cy5.5 | 1000 | 29F.1A12 | BioLegend | 135208 |
| LAG3 | BV650 | 100 | C9B7W | BioLegend | 125227 |
| TIM3 | BV785 | 100 | RMT3-23 | BioLegend | 119725 |
| CD19 | BUV661 | 800 | 1D3 | BD Biosciences | 612971 |
| NKp46 | BUV395 | 200 | 29A1.4 | Biolegend | 568630 |
| Live/Dead | IR-876 | 1000 | - | Invitrogen | 174497772 |

**Supplementary Table 5**

| **Supplementary Table S5: Table of Culture Media composition** | | |
| --- | --- | --- |
| **Component** | **RPMI 1640 GlutaMAX Ref 61870-010** | **DMEM high glucose + pyruvate Ref 41966-029** |
| **1. Amino-acid (mg/L)** |  |  |
| Glycine | 10 | 30 |
| L-Arginine / HCl | 200 | 84 |
| L-Asparagine | 50 | — |
| L-Aspartic acid | 20 | — |
| L-Cystine | 50 | 63 |
| L-Glutamine / derivado | 446 (Ala-Gln) | 580 (L-Gln) |
| L-Glutamic acid | 20 | — |
| L-Histidine | 15 | 42 |
| L-Hydroxyproline | 20 | — |
| L-Isoleucine | 50 | 105 |
| L-Leucine | 50 | 105 |
| L-Lysine | 40 | 146 |
| L-Methionine | 15 | 30 |
| L-Phenylalanine | 15 | 66 |
| L-Proline | 20 | — |
| L-Serine | 30 | 42 |
| L-Threonine | 20 | 95 |
| L-Tryptophan | 5 | 16 |
| L-Tyrosine | 20 | 72 |
| L-Valine | 20 | 94 |
| **2. Vitamins (mg/L)** |  |  |
| Biotin | 0,2 | — |
| Choline chloride | 3 | 4 |
| D-Calcium pantothenate | 0,25 | 4 |
| Folic acid | 1 | 4 |
| Niacinamide | 1 | 4 |
| Para-aminobenzoic acid | 1 | — |
| Pyridoxine HCl | 1 | 4 |
| Riboflavin | 0,2 | 0,4 |
| Thiamine HCl | 1 | 4 |
| Vitamin B12 | 0,005 | — |
| i-Inositol | 35 | 7,2 |
| **3. Inorganic salts (mg/L)** |  |  |
| Calcium nitrate / chloride | 100 | 264 |
| Ferric nitrate | — | 0,1 |
| Magnesium sulfate | 100 | 200 |
| Potassium chloride | 400 | 400 |
| Sodium bicarbonate | 2000 | 3700 |
| Sodium chloride | 6000 | 6400 |
| Sodium phosphate | 800 | 141 |
| **3. other componentes** |  |  |
| D-Glucose | 2000 | 4500 |
| Sodium pyruvate | — | 110 |
| Glutathione (reduced) | 1 | — |
| Phenol red | 5 | 15 |

**Supplementary Figures**

**Supplementary Figure 1**

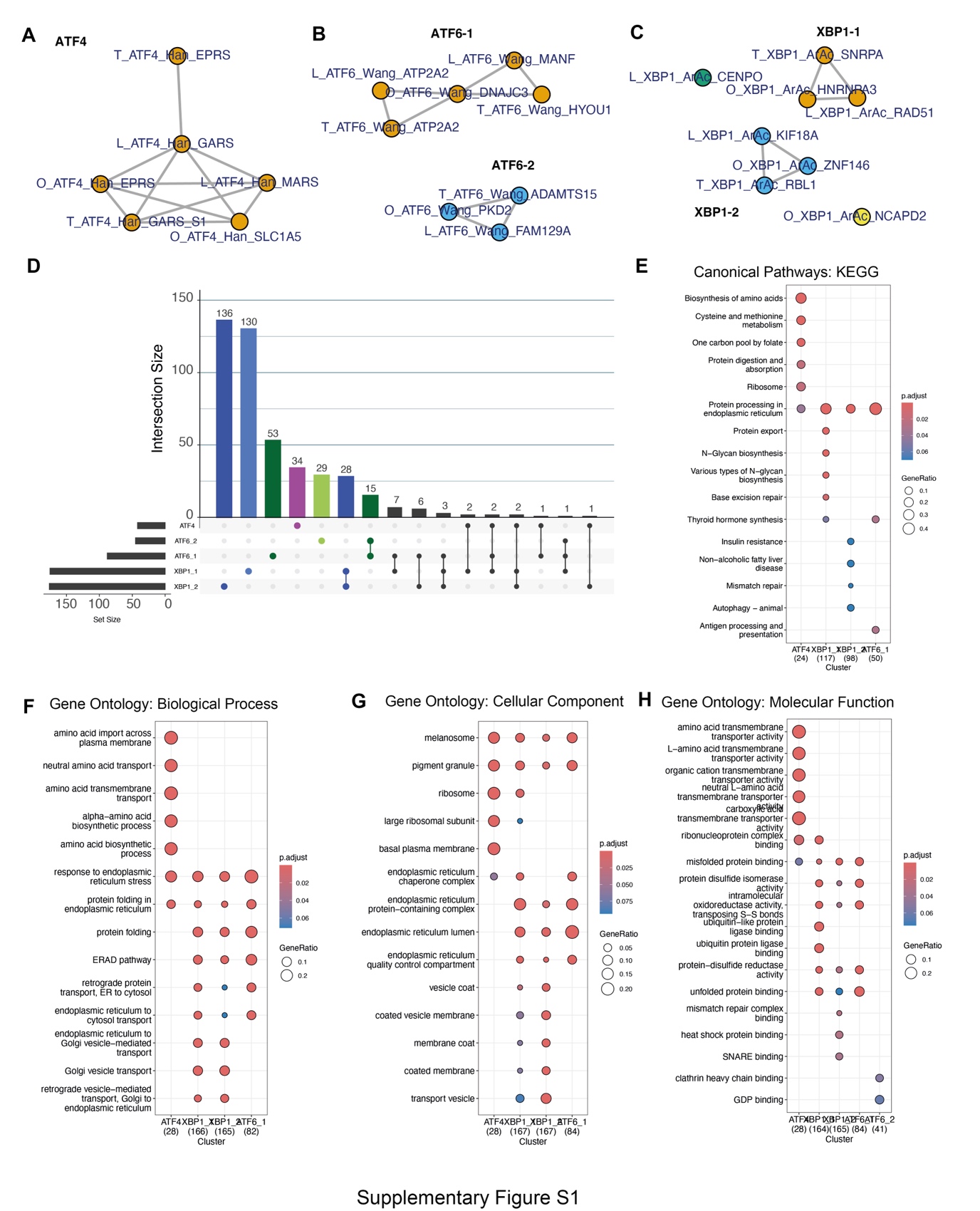

**Supplementary Figure 1. Functional characterization and overlap analysis of the gene signatures of ATF4, ATF6-1, and ATF6-2 and XBP1-1 and XBP1-2**

(A-C) Network diagrams showing the connectivity and relationships among the individual components of the ATF4, ATF6-1, and ATF6-2 and XBP1-1 and XBP1-2 gene signatures. Nodes represent specific signatures, and lines indicate significant coexpresión of signature scores. (D) UpSet plot showing the intersection and size of gene sets among the five ER stress signatures. The lower horizontal bars indicate the total size of each signature (Set Size), while the upper vertical bars quantify the genes shared among the combinations indicated by the connecting dots and lines (Intersection Size). (E-H) Functional enrichment analysis for the gene groups of each signature. Significant terms for the canonical KEGG pathways are presented, as well as the Gene Ontology (GO) categories: Biological Processes, Cellular Components, and Molecular Functions.

**Supplementary Figure 2**

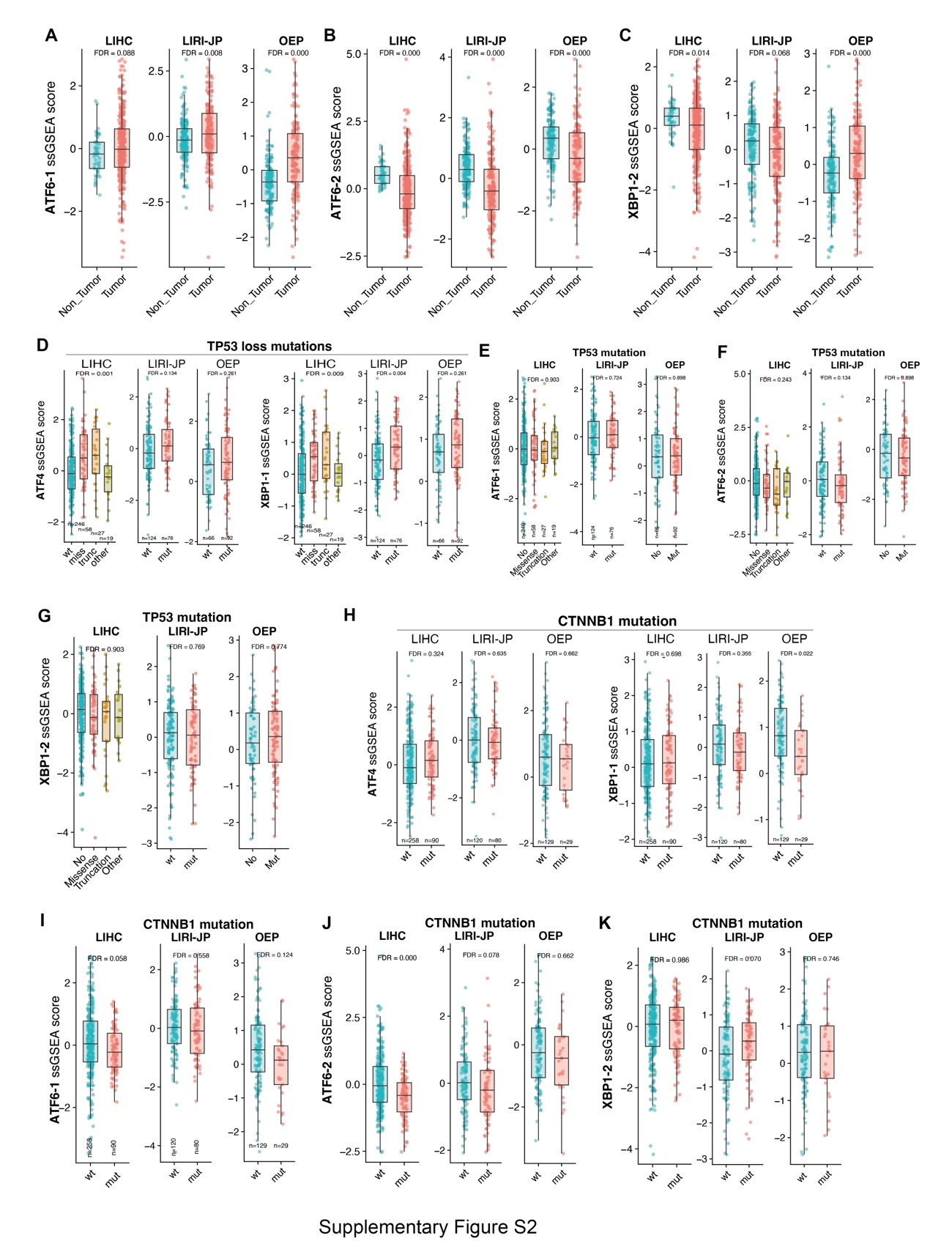

**Supplementary Figure 2. XBP1-2 and ATF6 signature scores were not increased in tumors.** (A-C) Comparison of single-sample gene set enrichment analysis (ssGSEA) scores for the ATF6-1, ATF6-2, and XBP1-2 signatures between non-tumor and tumor tissue. Data are shown for three independent cohorts: LIHC, LIRI-JP, and OEP. (D) ssGSEA scores for ATF4 and XBP1-1 categorized by *TP53* loss-of-function mutation status. (E-G) Scores for the ATF6-1, ATF6-2, and XBP1-2 signatures in relation to the presence or absence of TP53 mutations. (H-K) Association of ER stress signatures with *CTNNB1* (β-catenin) mutational status. Distribution of ssGSEA scores for ATF4 and XBP1-1, ATF6-1, ATF6-2 and XBP1-2 comparing tumors with wild-type (wt) versus mutated (mut) genotypes for *CTNNB1*.

**Supplementary Figure 3**

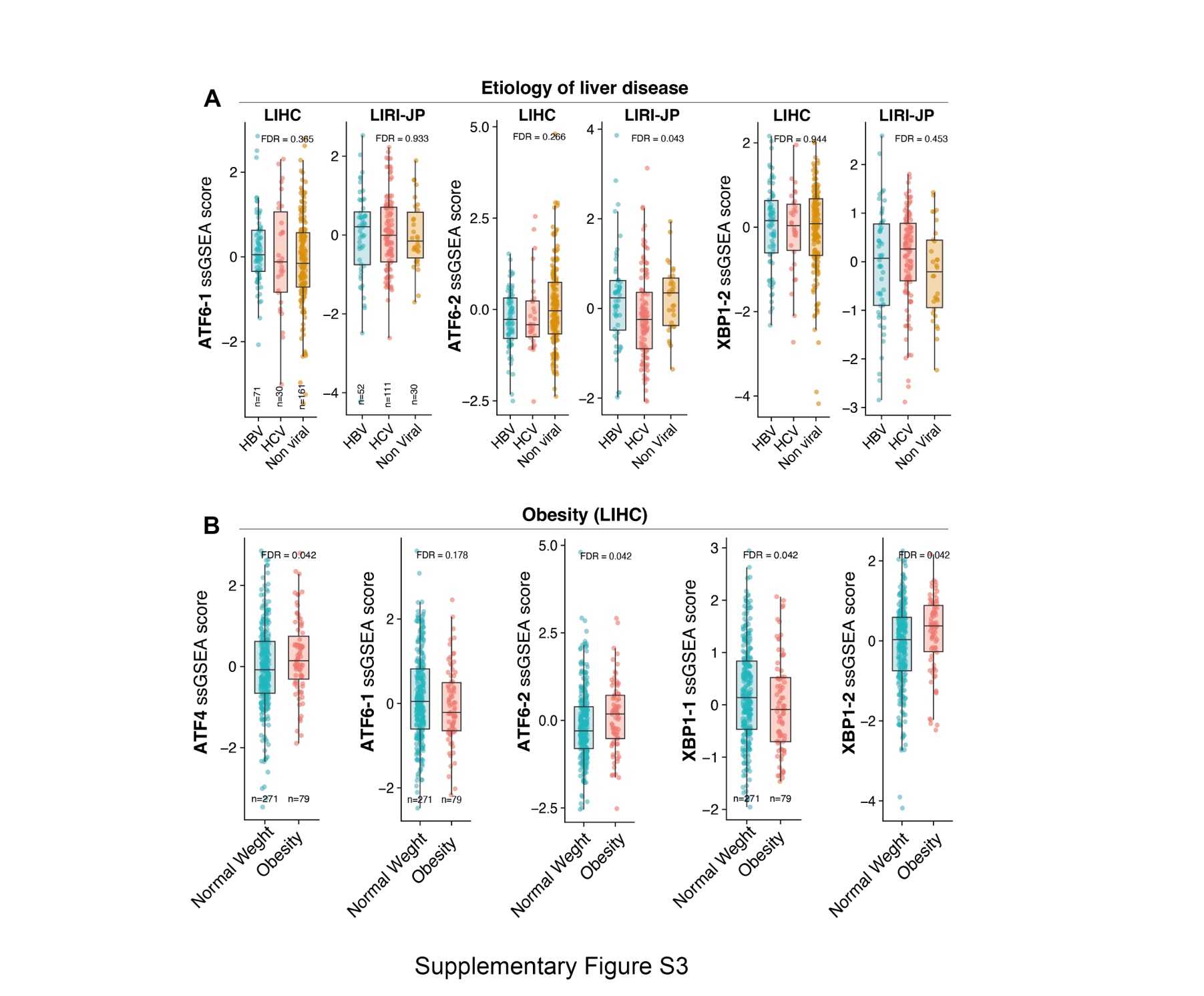

**Supplementary Figure 3. LIHC and LIRI cohorts, none of the 5 signatures were consistently associated with the etiology of the liver disease.** (A) Single-sample gene set enrichment analysis (ssGSEA score) for gene signatures ATF6-1, ATF6-2, and XBP1-2 in the LIHC and LIRI-JP cohorts, categorized by the etiology of the chronic liver disease: hepatitis B virus (HBV), hepatitis C virus (HCV), and non-viral etiology. (B) Comparison of ssGSEA scores for the signatures ATF4, ATF6-1, ATF6-2, XBP1-1, and XBP1-2 between normal-weight and obese patients within the LIHC cohort.

**Supplementary Figure 4**

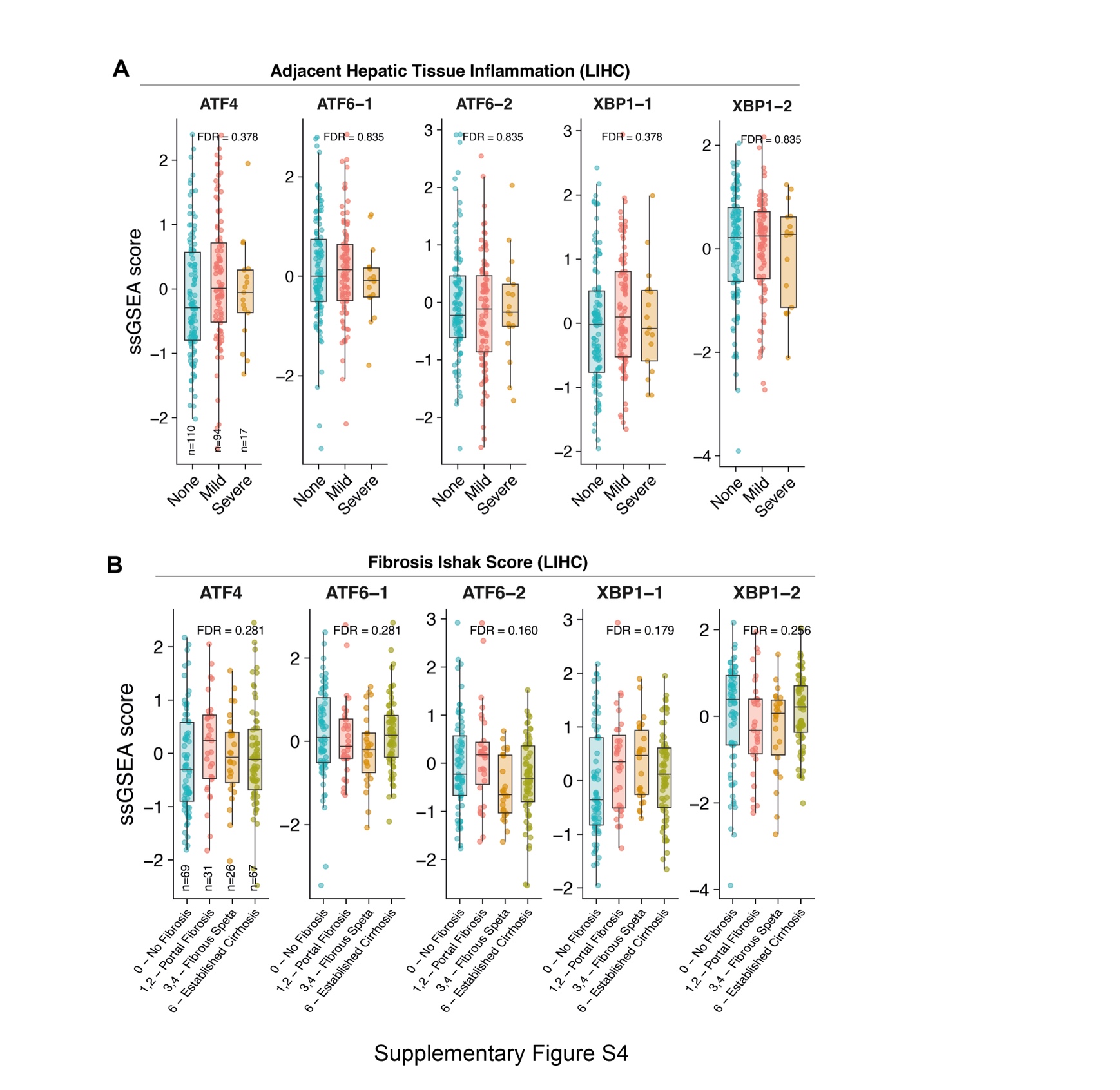

**Supplementary Figure 4. The UPR transcriptional activity in the tumor was not associated with the presence in the background liver of inflammation or fibrosis per Ishak score.** (A) Single-sample gene set enrichment analysis (ssGSEA score) for the gene signatures ATF4, ATF6-1, ATF6-2, XBP1-1, and XBP1-2 in adjacent liver tissue, categorized according to the level of histological inflammation: None, Mild, and Severe. (B) ssGSEA scores of transcription factor signatures distributed according to the Ishak scale for fibrosis: 0 (no fibrosis), 1–2 (portal fibrosis), 3–4 (fibrous septa), and 6 (established cirrhosis).

**Supplementary Figure 5**

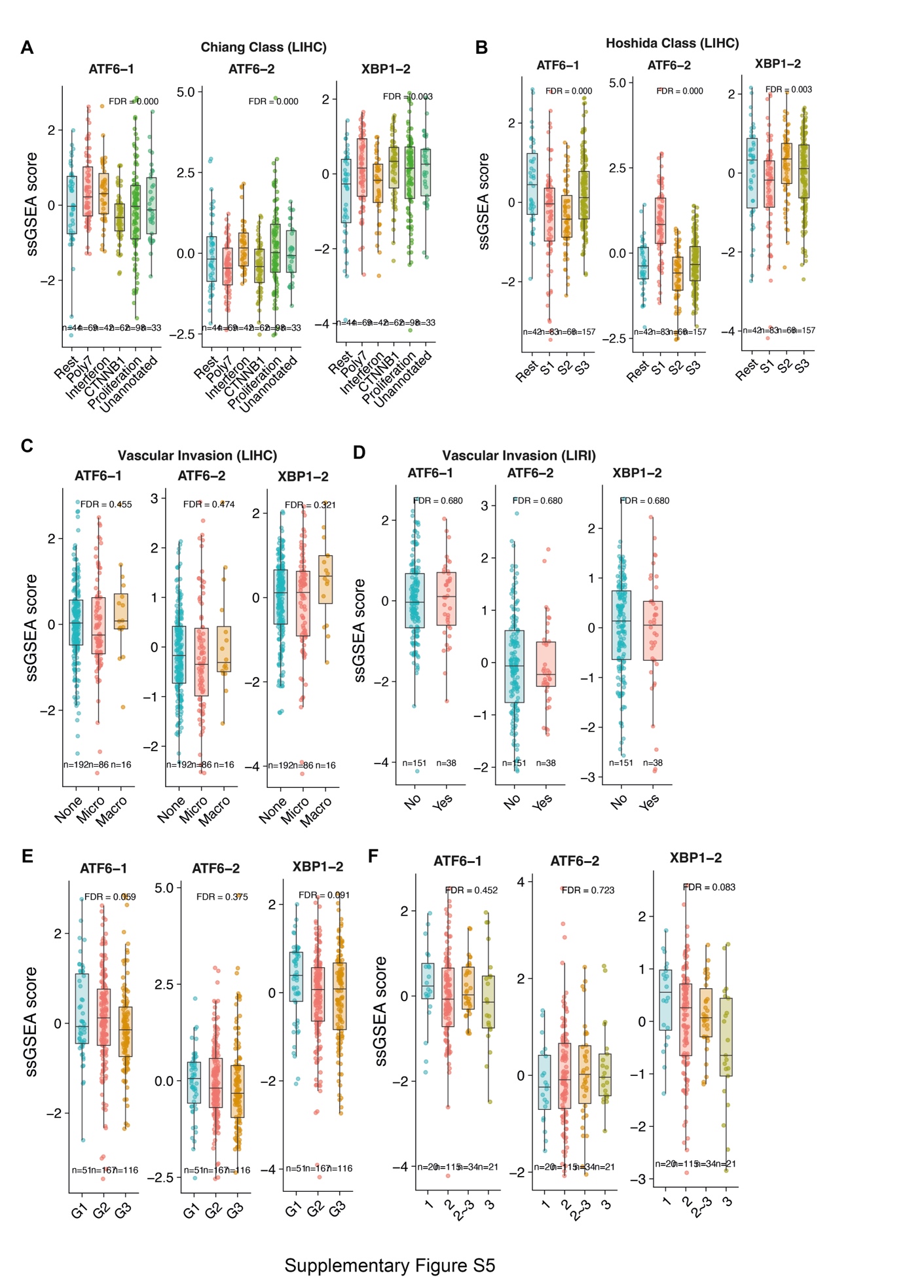

**Supplementary Figure 5. ATF6-1, ATF6-2 and XBP1-2 signatures, show no association with transcriptomic subclasses of Chiang or Hoshida, vascular invasion and differentiation tumors.** (A-B) ssGSEA scores for the ATF6-1, ATF6-2, and XBP1-2 signatures categorized by the Chiang and Hoshida molecular classifications in the LIHC cohort. (C-D) Comparison of ssGSEA scores of the transcription factor signatures in relation to vascular invasion in the LIHC cohorts, categorized as null (None), microvascular (Micro), and macrovascular (Macro), and LIRI, categorized in a binary fashion. (E-F) ER signature scores stratified by tumor histological grade in two independent cohorts, represented on a scale of 1 to 3.

**Supplementary Figure 6**

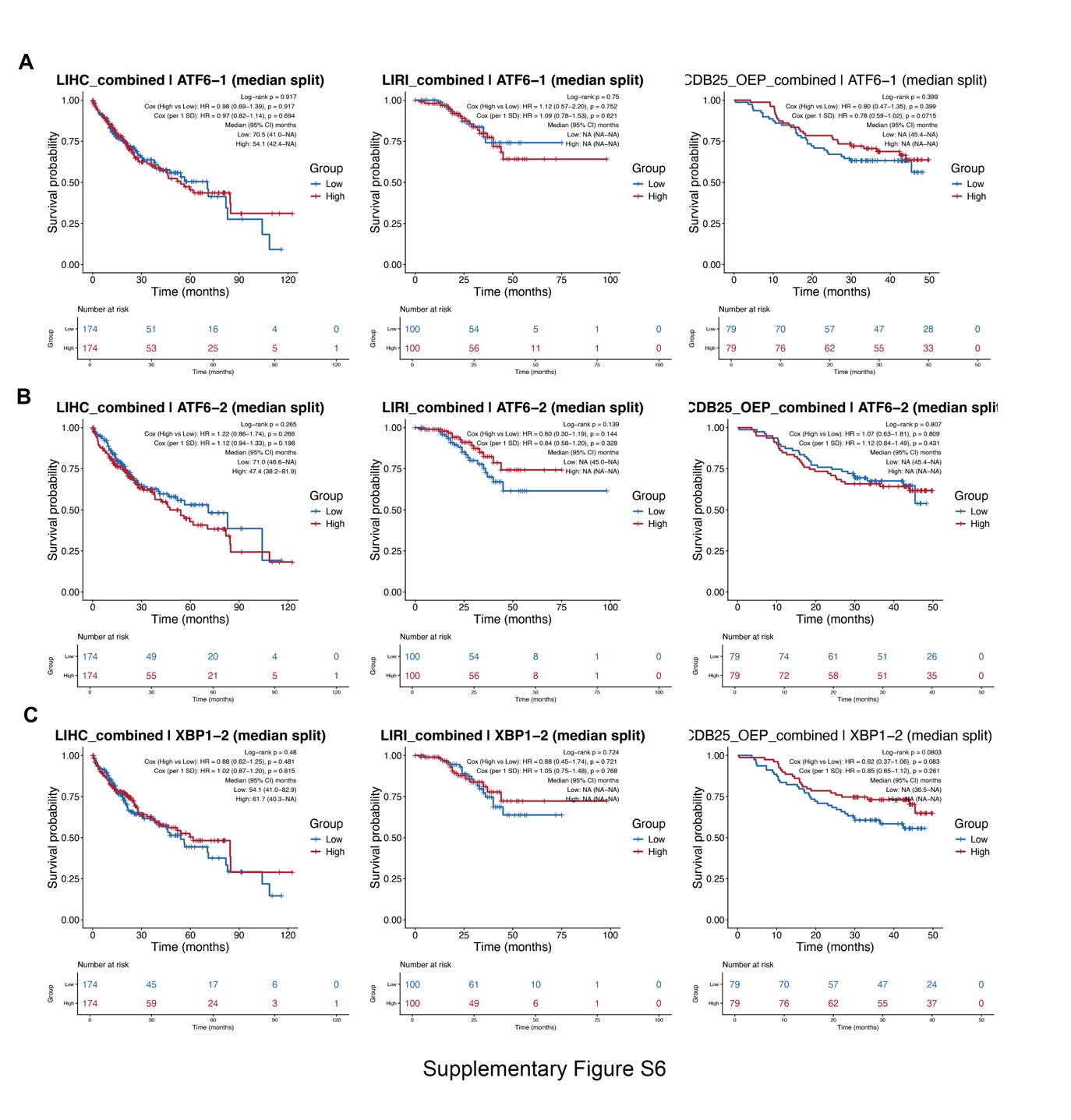

**Supplementary Figure 6. XBP1-2, ATF6-1 and 2 marked a group of patients with different prognosis.** (A-C) Overall survival Kaplan Meier curves for the gene signatures ATF6-1 (A), ATF6-2 (B), and XBP1-2 (C) in three independent cohorts (LIHC, LIRI, and CDB25_OEP). Patients were stratified into high (High, red) and low (Low, blue) expression groups by splitting by the median. Data of Log Rank (p value) and Cox Regression analyses (Hazard Ratio, HR and p value), and median survival for each group are presented on each plot, top right. NA stands for “not reached”.

**Supplementary Figure 7**

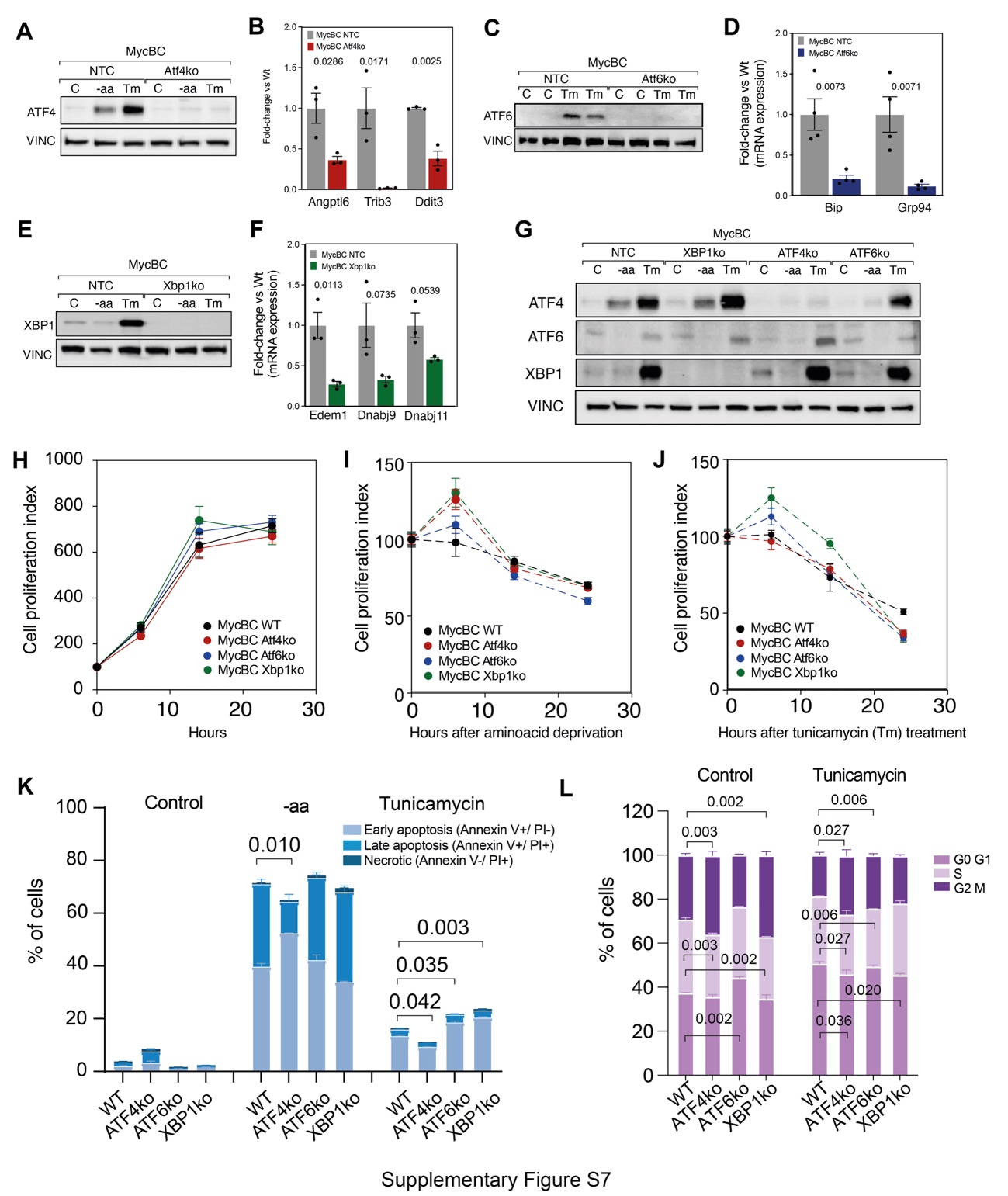

**Supplementary Figure 7. Atf4, Atf6, and Xbp1 knockout reveal limited effects on proliferation in vitro (**A-F) Knockout verification of UPR transcription factors by immunoblot analysis and RT-PCR. G) compensatory effects across UPR pathways. (H-J) Studies of cell proliferation index, (K) apoptosis and (L) cell cycle. Every assay was made under basal conditions (C) and stress conditions: amino acid deprivation (-aa) and N-glycosylation inhibition by tunicamycin (Tm). Data are shown as mean ± standard error of the mean (SEM) with individual biological replicates of three independent experiments. Statistical significance was assessed using two-way ANOVA followed by Dunnett’s multiple comparisons test comparing each Ko to NTC

**Supplementary Figure 8**

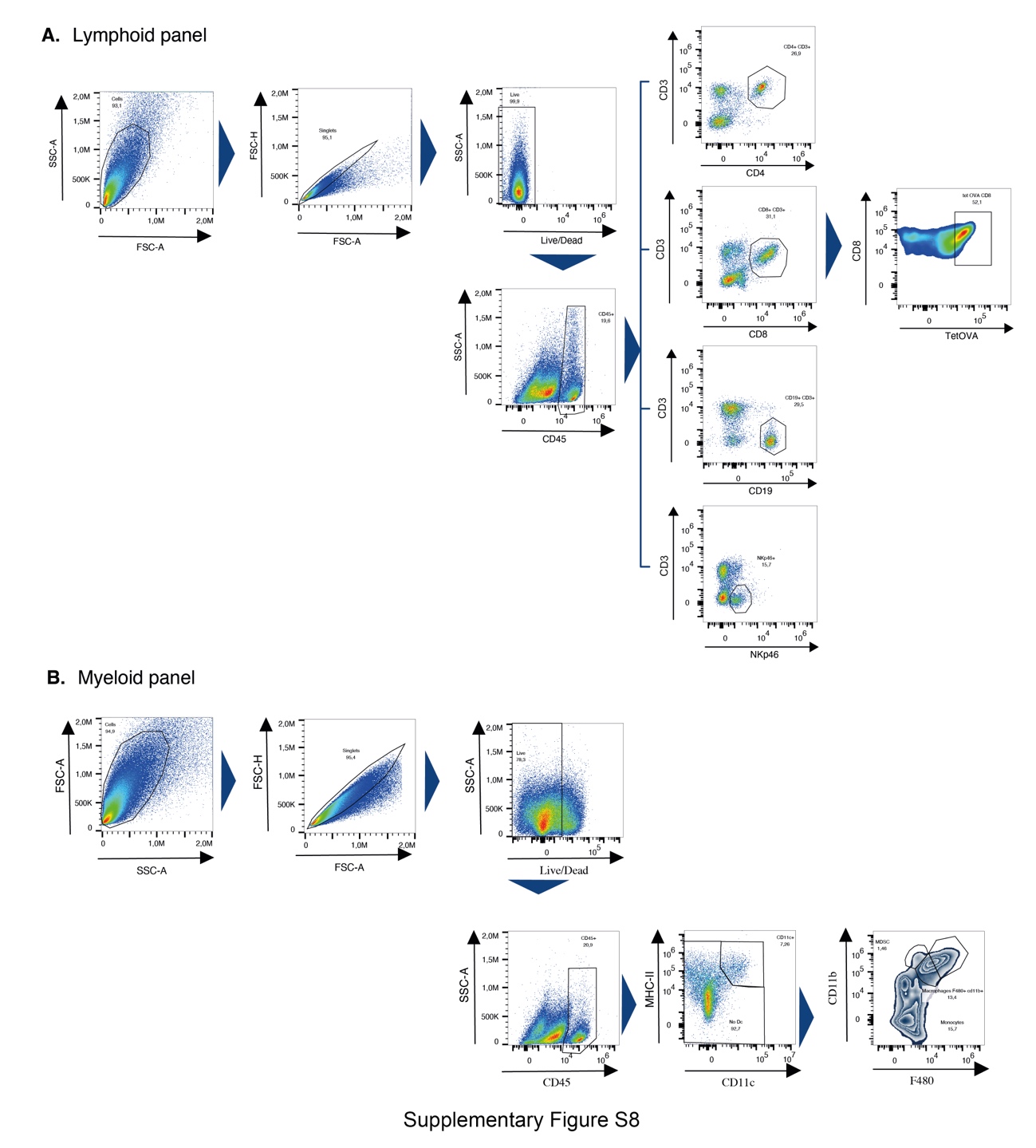

**Supplementary Fig 8. Gating strategy for the identification and study of immune cells in subcutaneus HCC MycBC tumors.** tumors were obtained from mice after 13 days of tumor inoculation. After digestion and homogenization, cells were stained with antibodies to identify lymphoid and Myeloid panel. Different were defined according to the gating strategy shown

**REFERENCES**

1.	Lu Y, Yang A, Quan C, Pan Y, Zhang H, Li Y, et al. A single-cell atlas of the multicellular ecosystem of primary and metastatic hepatocellular carcinoma. Nat Commun. 2022;13(1):4594.

2.	Ruiz de Galarreta M, Bresnahan E, Molina-Sanchez P, Lindblad KE, Maier B, Sia D, et al. beta-Catenin Activation Promotes Immune Escape and Resistance to Anti-PD-1 Therapy in Hepatocellular Carcinoma. Cancer Discov. 2019;9(8):1124-41.
